# Clone decomposition based on mutation signatures provides novel insights into mutational processes

**DOI:** 10.1101/2021.05.08.443215

**Authors:** Taro Matsutani, Michiaki Hamada

## Abstract

Intra-tumor heterogeneity is a phenomenon in which mutation profiles differ from cell to cell within the same tumor and is observed in almost all tumors. Understanding intra-tumor heterogeneity is essential from the clinical perspective. Numerous methods have been developed to predict this phenomenon based on variant allele frequency. Among the methods, CloneSig models the variant allele frequency and mutation signatures simultaneously and provides an accurate clone decomposition. However, this method has limitations in terms of clone number selection and modeling. We propose SigTracer, a novel hierarchical Bayesian approach for analyzing intra-tumor heterogeneity based on mutation signatures to tackle these issues. We show that SigTracer predicts more reasonable clone decompositions than the existing methods that use artificial data that mimic cancer genomes. We applied SigTracer to whole-genome sequences of blood cancer samples. The results were consistent with past findings that single base substitutions caused by a specific signature (previously reported as SBS9) related to the activation-induced cytidine deaminase intensively lie within immunoglobulin-coding regions for chronic lymphocytic leukemia samples. Furthermore, we showed that this signature mutates regions responsible for cell-cell adhesion. Accurate assignments of mutations to signatures by SigTracer can provide novel insights into signature origins and mutational processes.

## Introduction

Intra-tumor heterogeneity (ITH) is a phenomenon in which the mutation profiles differ from cell to cell within the same tumor and is observed in almost all tumors. In clinical practice (especially for treatment strategies), understanding heterogeneity is an important task because cell populations with heterogeneous genetic profiles make it challenging to determine which drugs are effective for a particular tumor [1]. In addition, heterogeneity represents how tumors have evolved. Hence, the accurate estimation of heterogeneity is essential to elucidate cancer dynamics. Multiregion sampling and single-cell DNA sequencing are effective in estimating the heterogeneity of a single tumor because they directly provide region-by-region and cell-by-cell mutational profiles. However, because of the high cost, low sequencing depth, and small amount of data acquired, *bulk* sequencing data are used in numerous cases for comprehensive analysis instead of single-cell sequencing.

ITH is mainly estimated via bulk sequencing by decomposing all mutations into temporally similar populations (called clones) based on the cancer cell fraction representing a hypothetical time axis. Numerous methods to predict ITH have been developed [2, 3, 4, 5, 6], and most depend on clustering mutations (i.e., decomposing into clones) by probabilistically modeling the variant allele frequency (VAF) for each sequenced mutation and calculating the cancer cell fraction (CCF) for each mutation by correcting the VAF using copy number aberrations (CNA) according to structural variants. However, the VAF of a single mutation is a noisy observation due to some technical limitations such as low sequencing depth, and it is difficult to accurately reconstruct the tumor evolution using only VAF.

When modeling mutations probabilistically, it is natural to focus on the cause of mutations, in other words, the mutational processes. In general, each mutational process leaves a specific fingerprint. This mutation spectrum can be formulated as a probabilistic distribution, called the *mutation signature* [7, 8]. For instance, the deamination of 5’-methylcytosine results in the characteristic single base substitution, N[C>T]G; hence, its mutation signature tends to have a higher proportion of these substitutions than others. The probabilistic distribution representing such a feature is registered in the COSMIC database as a signature (namely, SBS1), and more than 50 other signatures for single base substitutions have been reported. Several studies have suggested that each clone in the tumor is derived from different active signatures [9, 10, 11], and incorporating signatures to the model for estimating ITH was found to be useful. Abécassis, *et al* proposed CloneSig [12] that models mutation signatures and VAF simultaneously. CloneSig was found to outperform conventional methods of clone estimation with simulated mutation profiles that mimic whole-exome sequencing samples. While jointly modeling VAF and signatures, CloneSig considers that all mutations that accumulate in one tumor occur due to multiple signatures and each clone differs in the signature composition. For example, the clones with a strong SBS1 signature are expected to carry more N[C>T]G point mutations than other clones, as mentioned above. Therefore, mutation-clone matching can be achieved by following multiple clues including VAF and the substitution type and observing the surrounding bases, which improves the accuracy of clone estimation.

CloneSig has enabled many prospects for clone decomposition, but there are still some limitations. One of the drawbacks of CloneSig is the inaccurate model selection in terms of the number of clones using the Bayesian information criterion (BIC). BIC does not support rigorous validity of singular models [13] or mixture models, which have hidden variables. Another limitation of CloneSig is the instability of the estimation due to parameters with no prior distributions. In this study, we developed SigTracer, a hierarchical Bayesian extension of CloneSig that provides a method of valid clone number selection and more robust clone decomposition, which selects the clone number using the evidence lower bound (ELBO), the lower bound of the Kullback–Leibler (KL) divergence between the true distribution and the approximate posterior distribution. Besides, SigTracer prepares Dirichlet distributions as conjugate priors for the signature activity of each clone. This extension can be regarded as a generalization of the CloneSig model, and other studies on similar tasks such as signature extraction have already highlighted the effectiveness of Dirichlet priors [14, 15, 16]. Here, we aimed to show if SigTracer provides more reasonable clone estimations than CloneSig for artificial tumors.

Clone decomposition based on mutation signatures also has significant potential in terms of signature analysis. Although methods to estimate which signatures are active in a given tumor have been proposed[17, 18], they usually do not consider VAF. In other words, they infer which signature leads to a certain mutation from only its trinucleotide type and the mutational distribution of the signature. This can lead to incorrect assignment of mutations to signatures. If each clone in the same tumor has a different active signature, these methods may incorrectly presume that a signature that is not active in the clone carrying a certain mutation yielded that mutation. In fact, in the original paper reporting CloneSig, an example of a sarcoma patient was provided who had clones with differently dominant signatures within a single tumor, and other studies have also suggested the transition of signature activities in coordination with clones [9, 10, 11]. All these results indicate the effectiveness of considering VAF for the analysis of mutation signatures. We applied SigTracer to whole-genome sequences of blood cancer samples and reported the findings through accurate signature assignment.

## MATERIALS AND METHODS

### Overview of the SigTracer model and the generative process

Figure 1 shows the graphical model of SigTracer for a single tumor, and we have summarized all notations in Table S1 of the Supplementary Materials. The tumor contained a total of *N* point mutations, and for every mutation, we simultaneously modeled the mutation type *x*_*n*_(1 ≤ *n* ≤ *N*) and the number of reads overlapping *x*_*n*_. *B*_*n*_ and *D*_*n*_ indicate the number of reads with mutated and total alleles, respectively. We considered six types of single base substitutions and 16 different neighboring bases around the mutated base using the known SBS signature set; thus, *x*_*n*_ was a categorical variable, and it took *V* = 96 different values. Relatedly, assuming that the subset consisting of *K* active signatures in the tumor was known and each mutational distribution of the *k*-th signature was denoted by ***φ***_*k*_ ∈ ℝ^*V*^ for 1 ≤ *k* ≤ *K*, we easily estimated the subset by applying various fitting methods [17, 18] to the mutation set in advance. For the genomic locus in which each mutation existed, we assumed that the copy number in a cancer cell 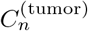, the copy number of a major allele in a cancer cell 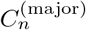, the copy number in a normal cell 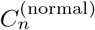 (these are collectively denoted as *C*_*n*_ in Figure 1), and the sample purity *P* were also known, and we must set these values in some way to predict CNA considering structural variants [19, 20].

**Figure 1:**
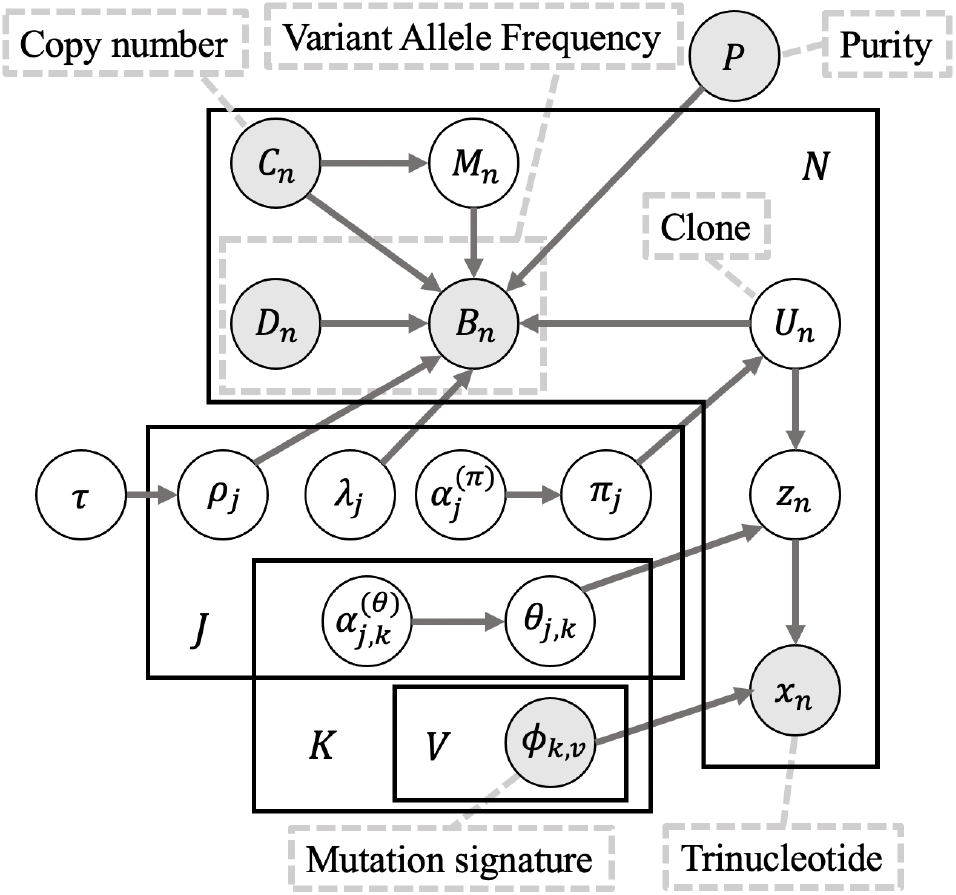
Graphical model of SigTracer for one tumor sample. This representation follows the plate notation, in which the variables shaded in black represent constants that can be observed in advance. Notations for all variables are explained in the main text and Table S1 of the Supplementary Materials.

For each mutation, the SigTracer model had three latent variables—*z*_*n*_, *U*_*n*_, and *M*_*n*_—indicating the signature via which the mutation occurred (a categorical variable with *K* types), the clone carrying that mutation (a categorical variable with *J* types where *J* is the number of clones), and the copy number of the mutation (*M*_*n*_ ∈ ℕ satisfying 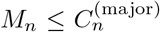), respectively. The clone *U*_*n*_ and signature *z*_*n*_ were generated using categorical distributions with parameters ***π*** ∈ ℝ^*J*^ and 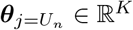, which followed prior Dirichlet distributions with ***α***^(*π*)^and 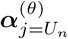. Note that each clone *j* had a different signature activity ***θ*** _*j*_. *M*_*n*_ was also modeled to follow a categorical distribution so that any possible natural number less than or equal to 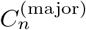 was equally sampled. Then, the number of reads, *B*_*n*_ and *D*_*n*_, was probabilistically modeled as follows: where

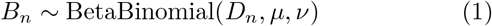

Where

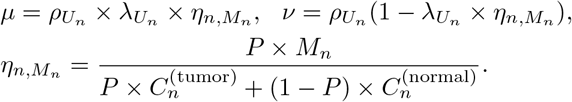

Here, *λ*_*j*_ and *ρ*_*j*_ are the CCF and overdispersion parameters for the *j*-th clone, respectively. BetaBinomial(*N, µ, ν*) is the probabilistic distribution of the number of successes in *N* trials, where success probability is sampled from a prior beta distribution with shape parameters *µ* and *ν. η*_*n,m*_ is the normalization term for the copy number of the mutation *n* in a sampled cell when *M*_*n*_ = *m* and the expected value of Beta(*µ, ν*) becomes 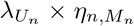. CCF, which is the VAF corrected for copy number, is defined as the proportion of sequenced cancer cells that contain mutations. *λ*_*j*_, which is the CCF for the *j*-th clone, shows when the clone was established (the larger the *λ*, the older the clone) under certain assumptions, including the infinite-site model.

In summary, the entire generative process is given as follows:

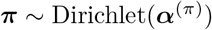

**for** each clone *j* = 1, …, *J* **do**

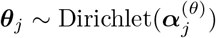

**for** each mutation *n* = 1, …, *N* **do**

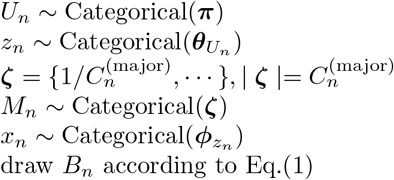

### Inference algorithm

#### Estimation of hidden variables and parameters

We predicted the responsibilities of the latent variables using the collapsed variational Bayesian (CVB) inference, and for marginalized parameters (i.e., ***π*** and ***θ***), we computed the estimates using the expected values obtained from the approximate posterior distributions and hyperparameters. *q*(***z, U***, ***M***), *q*(***π***), and *q*(***θ***) were the approximated posterior distributions of the latent variables and the corresponding parameters. Regarding latent variables, when we used the mean-field approximation, such as *q*(***z, U***, ***M***) ≈ *q*(***z***)*q*(***U***)*q*(***M***), preliminary experiments showed that the prediction accuracy was significantly worse. Therefore, while preserving the structure among the latent variables (i.e., joint posterior *q*(***z, U***, ***C, π, θ***) is factorized into 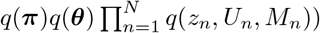, we derived the objective function, ELBO, as follows:

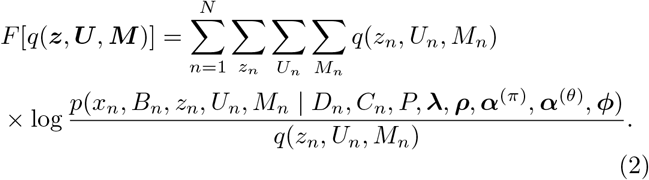

According to this objective function, we can obtain the updated formula for *q*(*z*_*n*_, *U*_*n*_, *M*_*n*_) which takes a stationary point to give the extreme value of *F* [*q*(***z, U***, ***M***)] as follows:

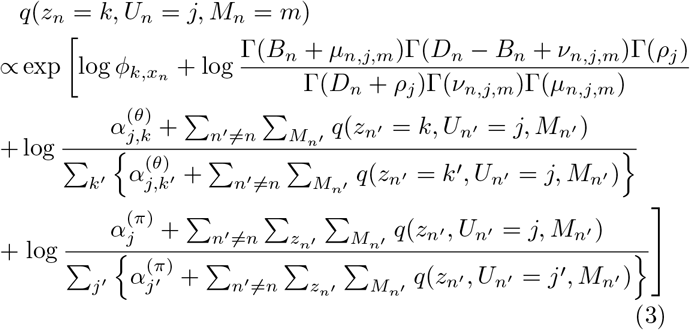

where Γ(·) denotes the gamma function and

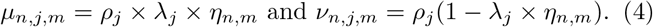

The detailed derivation is provided in Section 1 of the Supplementary Materials. For ***π*** and ***θ***, by taking the expected values with respect to *q*(*z*_*n*_, *U*_*n*_, *M*_*n*_) and hyper-parameters, we can estimate the following:

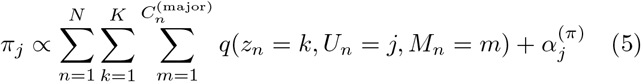

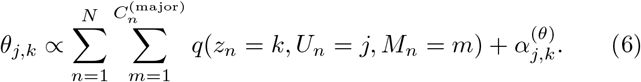

After estimating the responsibility and parameters, we calculated the expected CCF for each mutation as follows:

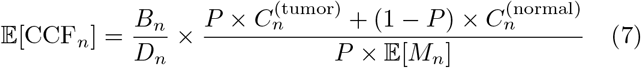

Where

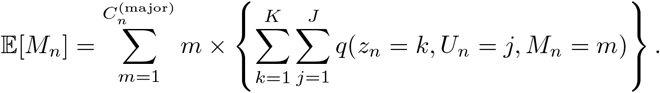

#### Estimation of hyper-parameters

We predicted the hyper-parameters, including ***α***^(*π*)^, ***α***^(*θ*)^, *λ*_*j*_ and *ρ*_*j*_, using fixed-point iterations to maximize ELBO. In fixed-point iterations, we first derived the lower bound of ELBO using the gamma function and used the stationary points to maximize it for each parameter. For the *ρ*_*j*_ update to control overdispersion, we employed the exponential distributions *p*(*ρ* | *τ*) = *τ* exp(*τρ*) as priors to achieve a stable estimation and confirmed that this modification prevented divergence of parameter learning. We have provided details of the updated formulas for all hyper-parameters in the Supplementary Materials (see Eqs. (S5)-(S12)).

#### Model selection using the variational Bayesian (VB) inference

To select a plausible number of clones, *J*, we used ELBO as the criterion for model selection in this framework. In CVB, although parameter estimation is possible using Eqs. (3)-(6), we could not explicitly calculate the ELBO value (as shown in Eq. (2)) because the approximate posteriors of ***π*** and ***θ*** were marginalized. Therefore, we used the VB method to derive the ELBO and performed tentative parameter estimation for model selection, which yielded the predicted number of clones for each tumor in advance of CVB. In the VB method, ELBO was formulated as shown in Eq. (S4) in Supplementary Materials, and we computed this value using the predicted parameters according to the update rules: Eqs. (S5)-(S7).

#### Different properties of SigTracer from CloneSig

In terms of modeling, one difference was that we prepared Dirichlet distributions as priors of signature activities for each clone (***θ***_*j*_) and the clone proportion (***π***). These categorical distribution parameters yielded *z*_*n*_ and *U*_*n*_, and CloneSig predicted these via an EM algorithm. Therefore, the update formula of the EM algorithm was equivalent to that of the VB method when the priors were Dirichlet distributions with all parameters set to 1.0. Another improvement in modeling was in terms of the setting for CCF overdispersion (*ρ*). CloneSig set the same value for all clones, whereas SigTracer controlled the overdispersion by each clone for the beta distribution, which is the prior of binomial distributions that determined *B*_*n*_ against *D*_*n*_.

Regarding an inference algorithm, SigTracer adopted ELBO as a criterion to select the number of clones instead of BIC used in CloneSig. In addition, we used fixed-point iteration to estimate hyper-parameters instead of the projected Newton method used by CloneSig.

#### Summary of the algorithm to infer parameters

The bottleneck of inference for both CVB and VB was calculated using *q*(***z, U***, ***M***), and the following time complexity 𝒪 (*NKJ* × max(***C***^(major)^)). To terminate CVB learning, we used the approximated ELBO. As described above, ELBO derived using CVB could not be calculated because the posteriors of ***π*** and ***θ*** were marginalized; hence, we approximated ELBO by substituting the expected value of ***π*** and ***θ*** into Eq. (S4). This value could not be used for model selection because it was not the objective of CVB, but it was used to confirm if training was saturated. Finally, we created an algorithm 1 to summarize the inference.

### Statistical test for measuring the relationship between mutations and signatures

We assumed that a particular somatic mutation drove tumor evolution and induced mutational processes of an individual signature, or conversely, a certain mutational process caused particular mutations. In that case, such mutations were likely to be concentrated in clones in which the relevant signature was highly active. Based on this idea, we implemented the following pipeline of statistical tests to measure the relevance between mutations and signatures at a genetic level.

First, for all mutations, we determined the clone *ĵ* to which

#### Algorithm 1 Parameter estimation of SigTracer

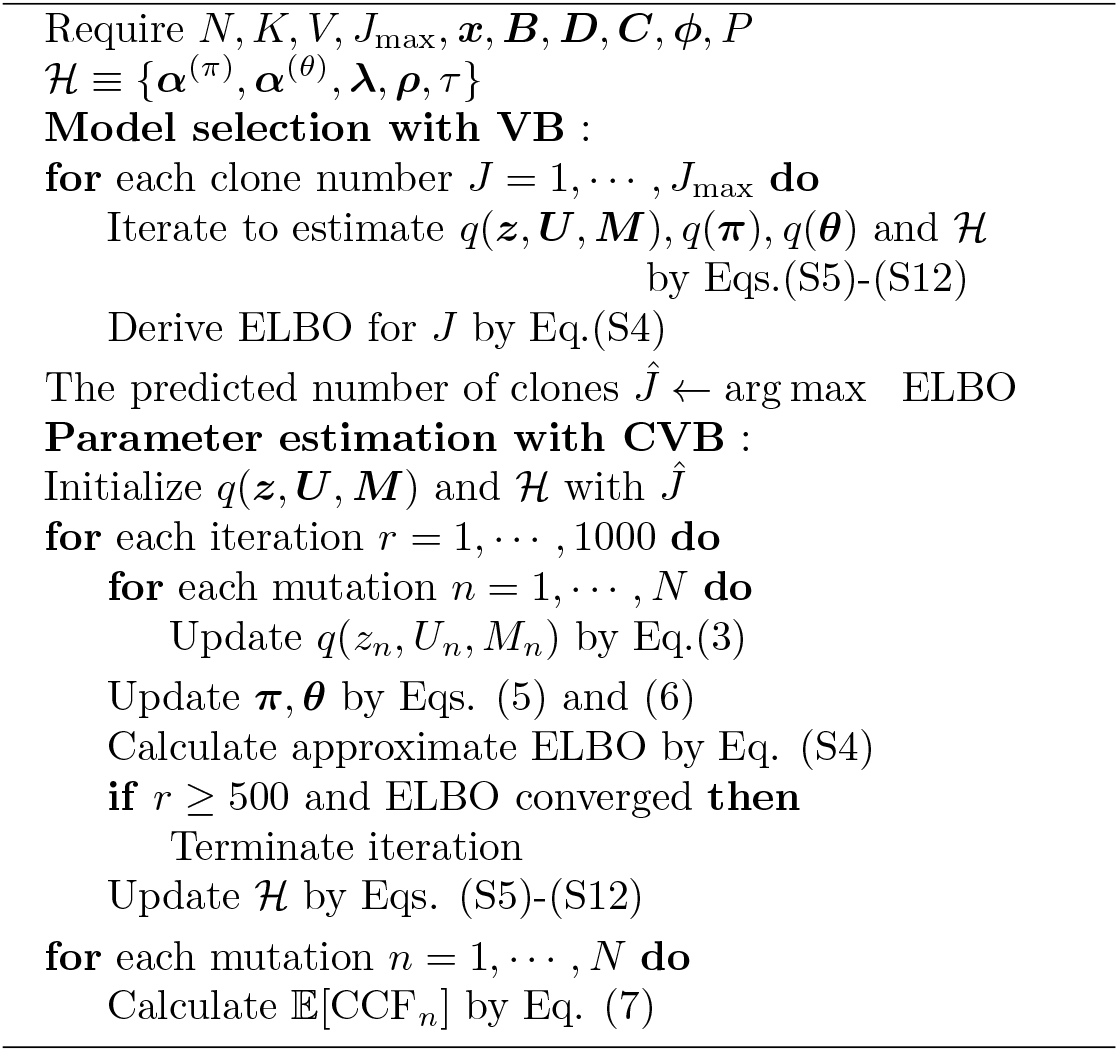

the mutation *n* belonged, as follows:

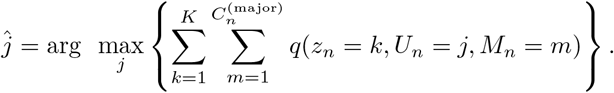

The next step was to determine the active signature *k* for each clone *j* based on whether or not the following threshold was satisfied:

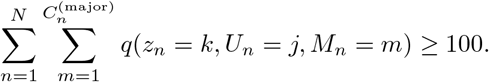

We divided all clones into active or inactive groups in terms of the signature *k* to be tested according to the above threshold, and we calculated the ratio of active group sizes for the signature *k* against all groups (denoted as *r*_*k*_) by adding the number of mutations in the clones belonging to each group.

If we knew where the mutations occurred at the genetic level, we estimated whether the clones carrying mutations on a certain gene were active or inactive for a particular signature *k* according to the above procedures. If there was no relationship between the gene of interest and the signature *k*, the mutations would be distributed into active and inactive groups according to the ratio *r*_*k*_. To test this null hypothesis, we performed a binomial test for all gene-signature combinations against a binomial distribution with a success probability of *r*_*k*_. In the actual implementation, we used scipy.stats.binom test with a significance level of *α* < 0.05 in Python.

### Datasets

#### Simulation data

To evaluate the performance of SigTracer, we artificially produced 14 datasets following the generative process in SigTracer. All datasets contained 100 samples, and we used 67 signatures registered in COSMIC ver3.1 as the reference mutational distributions. The datasets were divided into three categories: whole-genome sequencing (WGS) mimicking (low coverage and a large number of mutations); whole-exome sequencing (WES) mimicking (high coverage and a small number of mutations); and ideal (high coverage and a large number of mutations). Detailed information regarding how the datasets were produced is provided in Section 2 of the Supplementary Materials, and Table 1 summarizes the differences in essential parameters for each dataset.

**Table 1:**
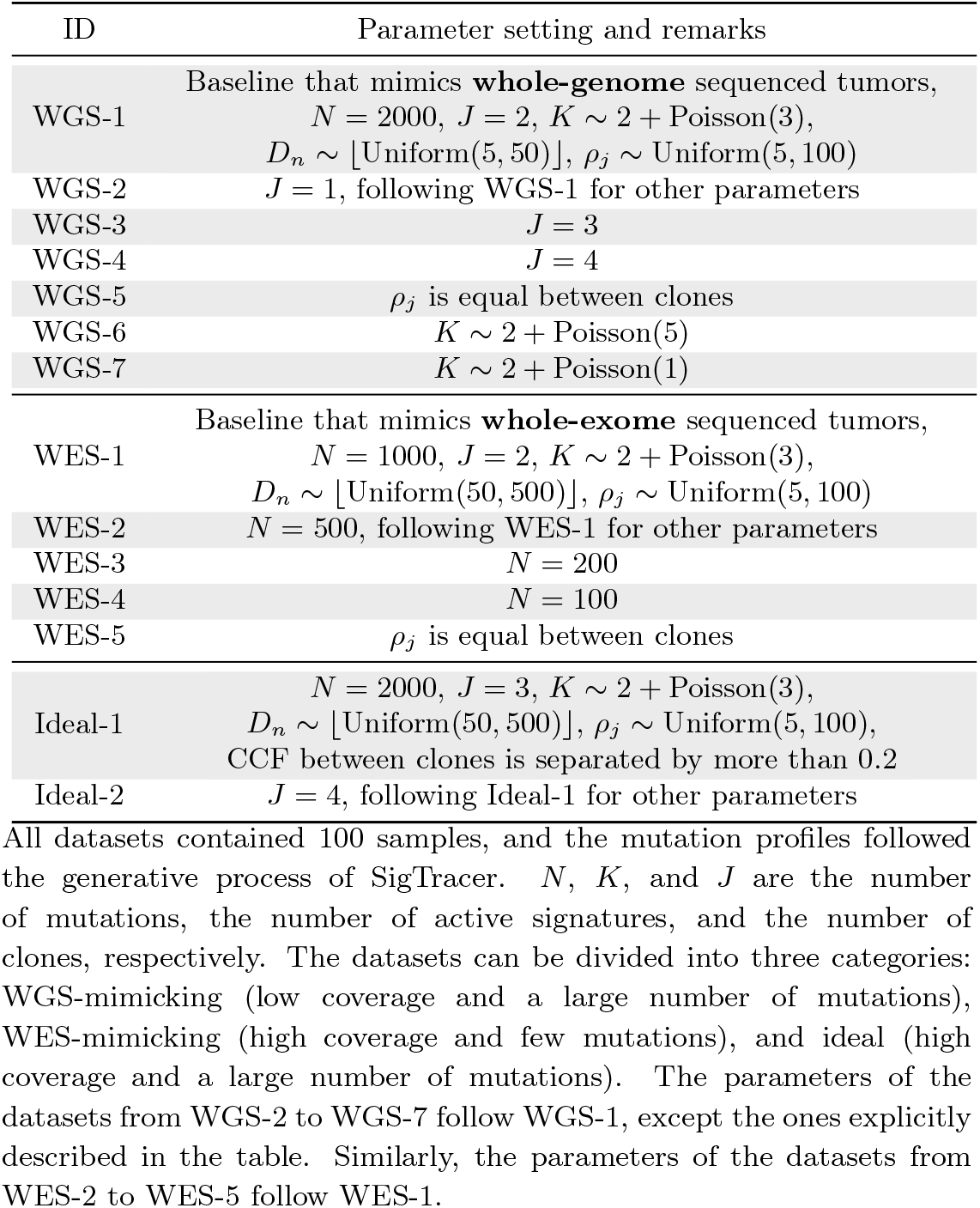
Artificial datasets and the setting of essential parameters.

### Application using blood cancer samples

As a proof of concept, we applied SigTracer to blood cancer samples of the Pan-Cancer Analysis of Whole Genomes (PCAWG) cohort. Tumors used in this study were subjected to whole-genome sequencing, and the tumor types were roughly divided into two categories: chronic lymphocytic leukemia (CLL) and B-cell non-Hodgkin lymphoma (BNHL), including Burkitt lymphoma, diffuse large B-cell lymphoma, follicular lymphoma, and Marginal lymphoma. All sources are available for download from the ICGC Data Portal: https://dcc.icgc.org/releases/PCAWG. We obtained 95 CLL samples and 100 BNHL samples. Similar to the simulation, we used COSMIC ver3.1 as the reference SBS signature set: https://cancer.sanger.ac.uk/cosmic/signatures/SBS/index.tt. For each tumor, previous studies using SigProfiler reported the types of active signatures [21]; four signatures, SBS1, SBS5, SBS9, and SBS40, were active in CLL, and 14 signatures, SBS1, SBS2, SBS3, SBS5, SBS6, SBS9, SBS13, SBS17a, SBS17b, SBS34, SBS36, SBS37, SBS40, and SBS56, were active in BNHL. This result is available in synapse.org ID syn11801889: https://www.synapse.org/#!Synapse:syn11804040, and we utilized them as the model input. Furthermore, the CNA and purity were estimated using several methods and are provided in the PCAWG database [22]; we also used these as the model input. In addition, when we applied the statistical test described above, we utilized the locus of all somatic mutations from mapping results uploaded on the PCAWG database, which were already annotated using Hugo symbols.

## RESULTS AND DISCUSSION

### Simulation experiments

#### Evaluation of model selection

To evaluate the model selection performance and compare SigTracer with CloneSig, we applied each method to the datasets presented in Table 1. We could easily identify active signatures in each sample using different fitting methods. Hence, we assumed that the type and number of signatures were known in this simulation. In addition, we elected *J*_min_ = 1 and *J*_max_ = 5 as the range of the clone number. Table 2 summarizes the results of model selection. Out of the 100 samples in each dataset, the bold characters indicate the number of samples for which the correct number of clones could be estimated using each method. We have presented the results of WES-2 to WES-5 in Table S2.

**Table 2:** The number of clones predicted with artificial mutation profiles.

| ID | Method | $J = 1$ | $J = 2$ | $J = 3$ | $J = 4$ | $J = 5$ |
| --- | --- | --- | --- | --- | --- | --- |
| WGS-1 | SigTracer | 16 | **81** | 3 | 0 | 0 |
|  | CloneSig | 1 | **25** | 42 | 23 | 8 |
| WGS-2 | SigTracer | **87** | 13 | 0 | 0 | 0 |
|  | CloneSig | **23** | 44 | 22 | 10 | 1 |
| WGS-3 | SigTracer | 6 | 72 | **21** | 1 | 0 |
|  | CloneSig | 0 | 12 | **39** | 35 | 14 |
| WGS-4 | SigTracer | 0 | 63 | 32 | **5** | 0 |
|  | CloneSig | 1 | 7 | 36 | **38** | 18 |
| WGS-5 | SigTracer | 23 | **77** | 0 | 0 | 0 |
|  | CloneSig | 0 | **30** | 49 | 20 | 1 |
| WGS-6 | SigTracer | 19 | **80** | 1 | 0 | 0 |
|  | CloneSig | 1 | **31** | 43 | 19 | 6 |
| WGS-7 | SigTracer | 7 | **91** | 2 | 0 | 0 |
|  | CloneSig | 0 | **25** | 49 | 21 | 5 |
| WES-1 | SigTracer | 12 | **82** | 6 | 0 | 0 |
|  | CloneSig | 1 | **21** | 27 | 17 | 34 |
| Ideal-1 | SigTracer | 0 | 20 | **78** | 2 | 0 |
|  | CloneSig | 0 | 0 | **21** | 30 | 49 |
| Ideal-2 | SigTracer | 0 | 8 | 50 | **39** | 3 |
|  | CloneSig | 0 | 0 | 7 | **29** | 64 |

Table 2 shows that SigTracer consistently estimated the correct number of clones compared to CloneSig for *J* ≤ 2. When a tumor included multiple clones, such as WGS-3 and WGS-4, the CloneSig estimation was more accurate than the SigTracer estimation. However, with the Ideal-1/2 datasets (high coverage and a high number of mutations), SigTracer estimated the correct number of clones in numerous samples. Even with an ideal dataset like Ideal-1/2, CloneSig predicted a larger number of clones than was true, indicating that BIC did not work correctly with the singular model. In addition, the CloneSig implementation adopted heuristics to compensate for the degrees of freedom in BIC, which might not be suitable for these cases. Notably, SigTracer exhibited better performance of model selection than CloneSig using the ideal datasets.

To investigate whether the tendency of SigTracer to estimate a small number of clones for the datasets with low coverage and many clones could be improved, we calculated the log-likelihood for each sample. Using artificial data, we could derive the “true” likelihood because we knew the correct latent variables (***z, U*** and ***M***) and true parameters (***λ*** and ***ρ***). Figure 2 shows the comparison between the log-likelihood based on the estimated parameters and the true log-likelihood for WGS-3, WGS-4, Ideal-1, and Ideal-2. Figure 2a and c show that the log-likelihood of SigTracer with fewer clones than the true number in WGS exceeded the log-likelihood with the true clone composition. In contrast, in the ideal datasets, the number of clones required by SigTracer was more than or equal to the true number to exceed the true log-likelihood in many samples. This result indicated that accurate clone number estimation in low-coverage data was challenging using the criteria based on the likelihood including BIC and ELBO.

**Figure 2:**
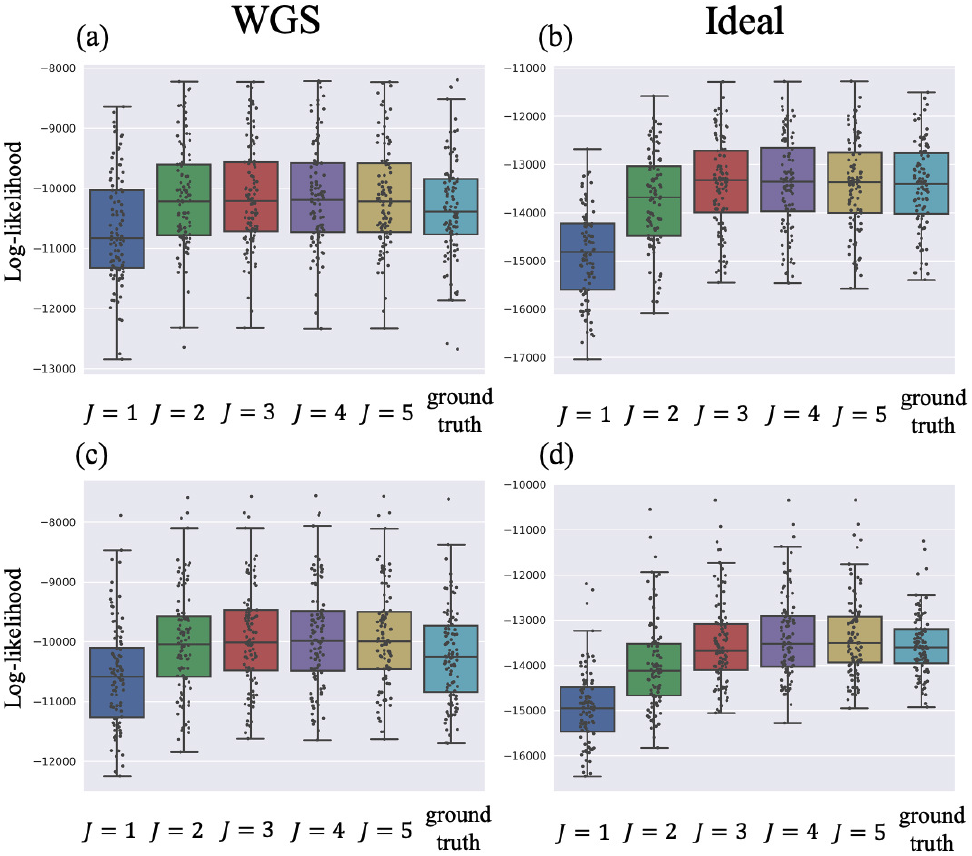
Log-likelihood with the estimated and ground-truth parameters in SigTracer. Each scatter plot shows the log-likelihood of each sample with the estimated and ground-truth parameters for *q*(***z, U***, ***M***), ***λ*** and ***ρ***. For the estimated parameters, box plots are drawn separately for the clone number. (a),(b),(c) and (d) differ in terms of the dataset, and they show the results of WGS-3, Ideal-1, WGS-4 and Ideal-2, respectively. The correct number of clones in (a) and (b) is *J* = 3, and that in (c) and (d) is *J* = 4.

Through these experiments, we highlighted the quantitative limitations of the current model for low-coverage data. However, its usefulness was still high, as evidenced by the fact that SigTracer could accurately estimate the clone number for numerous samples under an ideal setting. This method could become more critical as the number of high-coverage data will increase with the development of sequencing technologies in the future.

#### Evaluation of clone estimation

Next, we examined the accuracy of parameter estimations by SigTracer and CloneSig using the artificial datasets shown in Table 1. As measures of the estimation accuracy, we focused on whether they could correctly estimate CCF (***λ***) and the signature activity (***θ***) for each clone *j*. In this simulation, we provided the true number of clones and only evaluated the parameter estimation performance.

We denoted 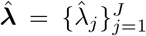 as the true CCF. For a single tumor, we defined the minimum value obtained by summing the CCF distance 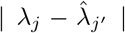 considering all possible combinations of the true and predicted clones as the evaluation criterion. For the signature activity by each clone, we denoted 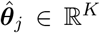 as the true activity for the *j*-th clone. We calculated the sum of the cosine distanced between ***θ***_*j*_ and 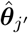 for all combinations, similar to the case of CCF, and used the minimum value among them as the evaluation criterion. Both criteria were desired to be small.

Figure 3 shows the results for a part of WGS dataset. We evaluated the results on an average for 100 samples included in each dataset. We have presented the results for other datasets in Figure S1. For CCF, except for WGS-4 with many clones and low coverage, SigTracer achieved comparable or better accuracy than CloneSig. In addition, the accuracy in terms of signature activity for each clone of SigTracer was comparable or better than that of CloneSig for all datasets. Finally, we have summarized the numerical statistics, including the mean and median values in Tables S3 and S4 for all simulation results.

**Figure 3:**
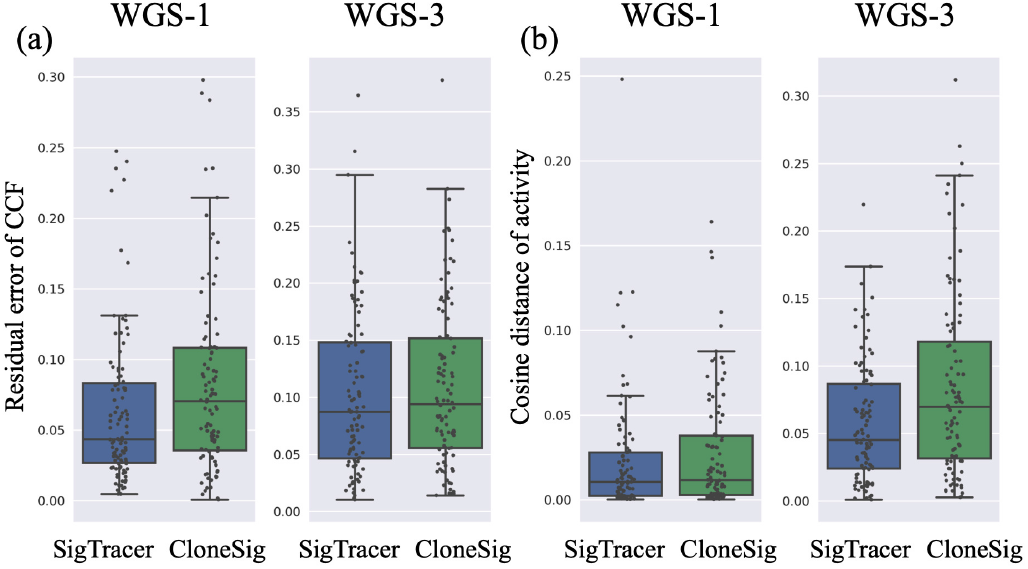
Accuracy of estimation for CCF and signature activity for each clone with the WGS-1/3 datasets. For the names of datasets, refer to Table 1. (**a**) and (**b**) show the residual error of the estimated CCF and the cosine distance of the estimated activity against the ground-truth values, respectively (the lower values are better). Blue box plots are the results of SigTracer, and green box plots are those of CloneSig. Each scatter plot shows the results of each sample included in each dataset. The results of other datasets are shown in Figure S1.

These improvements in accuracy were due to the differences in modeling as explained in MATERIALS AND METHODS. A clear example is the comparison between WES-1 and WES-5 shown in Figure S1. The only difference between these two datasets was whether the variance was set for each clone or not. SigTracer outperformed CloneSig in numerous samples in WES-1 in terms of CCF estimation, whereas there was almost no difference for WES-5. These differences were caused by the fact that SigTracer prepared the overdispersion parameters for each clone. Besides, in WGS with low coverage, SigTracer outperformed both CCF and the signature activity estimation for WGS-5 with the same variance between clones, suggesting that other modifications also contributed to the improvement.

#### Real data analysis with blood cancer samples

We applied SigTracer to CLL and BNHL samples described in the MATERIALS AND METHODS section. For model selection, the predicted numbers of clones are summarized in Figure S2. As observed in the simulation experiments, we must be aware that the predicted number of clones for WGS data might be lower than the actual number. Figure 4 shows an example of the output from SigTracer with one CLL sample, in which the expected value of CCF for mutation *n* was computed using Eq. (7) and was visualized by drawing a histogram from the responsibility of latent variables.

**Figure 4:**
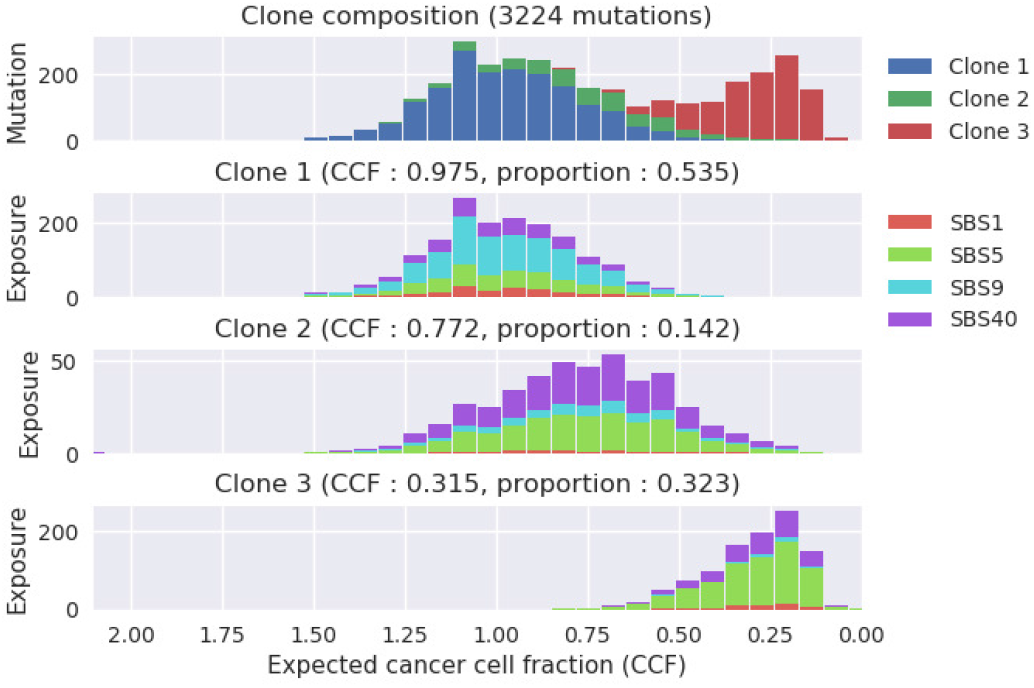
SigTracer output example with one CLL sample. The number of clones included in this tumor was estimated to be *J* = 3. The horizontal axis in all panels shows the expected cancer cell fraction. The top panel shows a histogram of all mutations with regard to the expected CCF, and it is color-coded to indicate the clone that the mutations belong to. Subsequent panels show histograms of mutations included in each clone, and they are color-coded to indicate which signature exposure is responsible.

Before we focus on the specifics, we provide some evidence to ensure the reliability of our results. First, we showed the validity of the SigTracer extension for preparing overdispersion parameters by each clone using the results obtained for CLL. Figure 5 summarizes the visualization of the reconstructed VAF distribution based on the estimated parameters when SigTracer was applied to a certain CLL sample with the number of clones, *J* = 3. Figure 5a shows the observed VAF (i.e., a histogram for *B*_*n*_*/D*_*n*_). Intuitively, when the VAF distribution was observed for a given number of clones, *J* = 3, it was desirable to decomposed the distribution into three elements with the expected VAF values of 0.1, 0.25, and 0.55. Figure 5b shows the result obtained when the variance of VAF was controlled by each clone (i.e., the model includes the extension of interest), which yielded an intuitively correct decomposition. In contrast, Figure 5c shows the result of SigTracer with the same value of overdispersions between clones; SigTracer predicted two clones with an expected VAF of almost 0.0. This tendency was not due to a simple difference in the initial values because the same results were obtained even after changing the initial values 10 times. The model with the same overdispersion parameters between clones could not capture the difference in the accumulation of mutations around the expected VAF. For instance, the red clone mutations in Figure 5b intensively occurred when the VAF was approximately 0.1, and the variance was smaller than those of the other two clones. However, in Figure 5c, this clone is shown to be split into two clones to adjust the variance between clones. Similar to this sample, we observed the effectiveness of the SigTracer extension in a number of CLL/BNHL samples.

**Figure 5:**
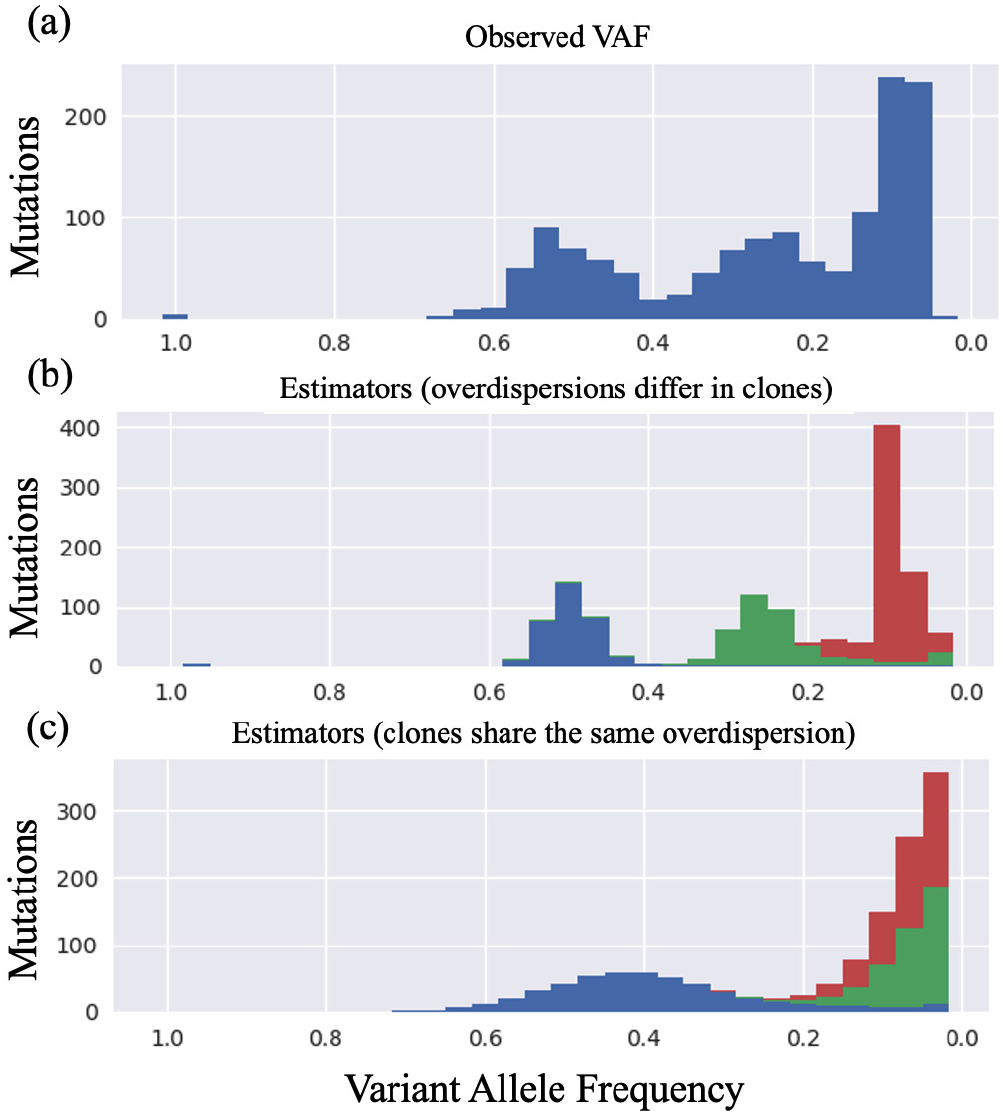
Comparison of the observed VAF in a certain CLL sample and the expected VAF based on the estimators of Sig-Tracer. The horizontal axis shows the VAF, and all panels show histograms for all mutations with respect to VAF. (**a**) is based on the observed VAF (i.e., a histogram with regard to *B*_*n*_*/D*_*n*_). (**b**) and (**c**) are histograms based on the estimated parameters under the observation of (**a**). In (**b**), SigTracer prepared overdispersion parameters for each clone (*J* = 3), and the figure is based on the estimators under those settings. In contrast, (**c**) is drawn based on the estimators obtained when SigTracer shares the same overdispersion across clones.

Next, we provide quantitative evidence that the clone decomposition by SigTracer for actual data was reliable. Because all signature-based methods include unsupervised learning and we cannot find true parameters, rigorously verifying the correctness of the estimated results for real data is not easy. However, based on the estimated parameters, we could quantify how accurately the model represents original data. We defined a measure called the “reconstruction rate: *RR*” to evaluate how many observed variables could be reconstructed. This measure was calculated by each sample. Two types of *RR*: *RR*_mutation_ were used to indicate how well the model reconstructed the mutation type (the *V*-dimensional categorical distribution) and *RR*_VAF_ to show how well the model reconstructed the VAF distribution (the beta mixture distribution); detailed definitions are provided in Section 4 of the Supplementary Materials. For instance, the *RR*_VAF_ calculation included quantifying the overlapping histograms in Figure 5a and b. By comparing the *RR* values calculated from the estimators using artificial and real data, we can see whether SigTracer was suitable for real data. In other words, if the *RR* value for the simulated data was close to that for real data, we could indirectly state that the real data could be represented by SigTracer. Table S5 summarizes the *RR* values calculated from all experiments. Although the *RR* values of real data (CLL and BNHL) tended to be lower than those of the simulated data that followed the generative process of SigTracer completely, we found the results of this study to be reliable to some extent. In summary, SigTracer is effective for not only artificial data but also for actual data.

#### SBS9 is initially active in CLL

SBS9 is the signature related with activation-induced cytidine deaminase (AID) or polymerase eta working in association with AID [8, 23]. The estimated CCF of the clones with high SBS9 activity tended to be close to 1.0 for most cases among the CLL samples, as shown in Figure 4. To quantify this trend, we divided the predicted clones into two categories: *primary clones* and *subclones*. We defined primary clone as clones with the largest size (*π*_*j*_) among clones with a CCF higher than 0.95 in one tumor. If no clone with a CCF higher than 0.95 existed, we defined the clone with the highest CCF in that tumor as the primary clone. Subclones were all clones except the primary clone. When we calculated which of the two types of clones was more likely to carry the mutations attributed to each signature from the estimated responsibility, we found that SBS9 was particularly active in the primary clone in CLL compared with other signatures (Figure 6a). This result was consistent with that of a previous report [22]. In contrast, this tendency was not observed in BNHL samples, and SBS9 was active in both primary and subclones (Figure 6b); hence, it remains to be elucidated what caused such a difference between the two types of blood cancer samples.

**Figure 6:**
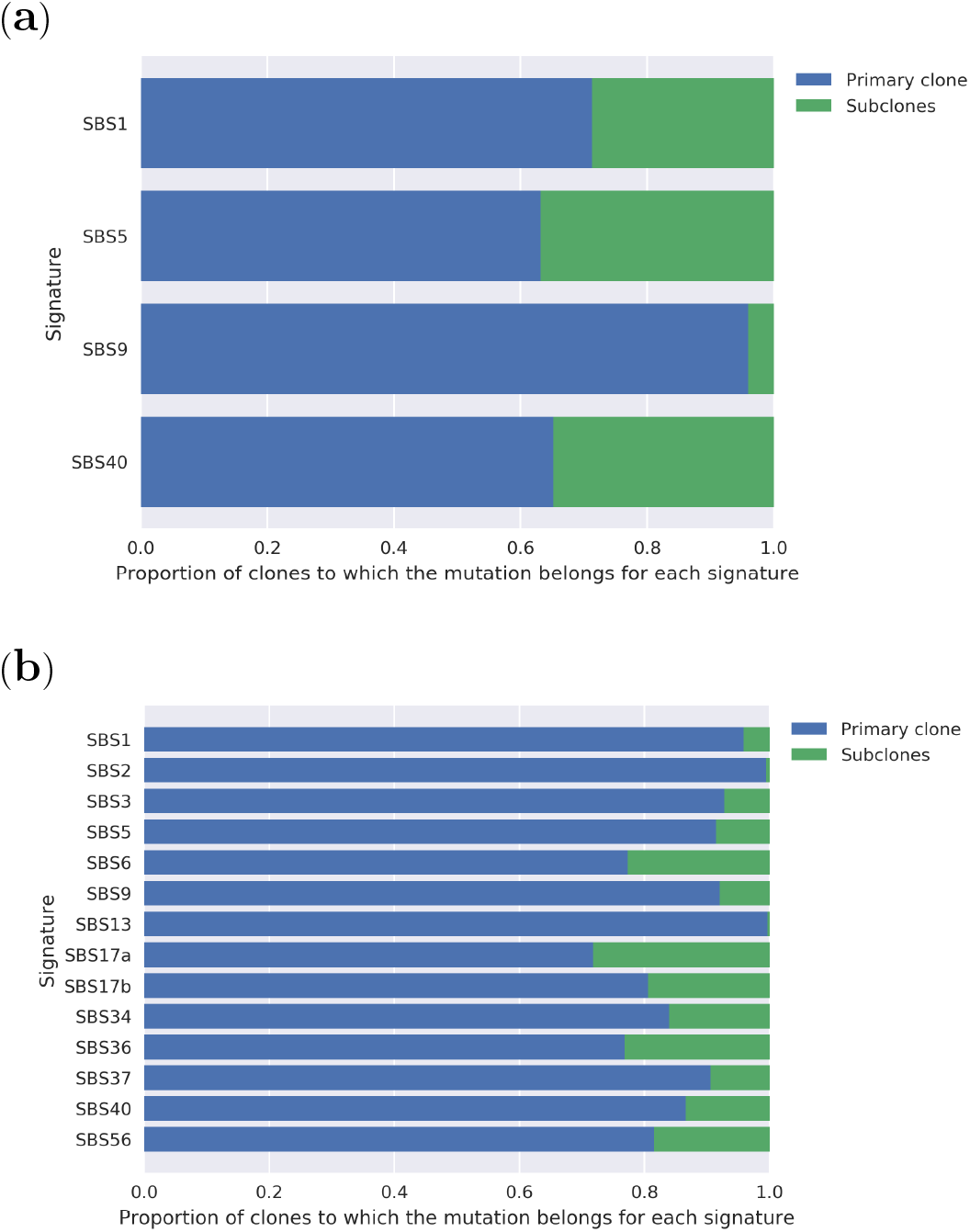
Types of clones that are likely to carry the mutations contributed by each signature. (**a**) and (**b**) show the results of CLL and BNHL, respectively. The clones were divided into two groups—primary and subclones—according to the procedure described in the text. This figure summarizes which signature-derived mutations they have. In CLL, the mutations in subclones contain a few SBS9-dreived ones (cf. the third bar in panel (a)).

#### Somatic mutations strongly associated with each signature

Using the statistical test pipeline described in the MATERIALS AND METHODS, we attempted to comprehensively identify mutations associated with each signature in CLL and BNHL samples at the genetic level. First, as a proof of concept, we have summarized the list of genes that were significantly associated with SBS9 in CLL samples in Table 3. We adopted the multiple test correction with familywise error rate (FWER) to avoid false positives. At the first glance, we observed that mutations were concentrated in the IG@@ region coding immunoglobulin. AID, the mutational process of SBS9, is required for class-switching of immunoglobulin [24]; thus, it is reasonable that mutations from SBS9 are concentrated in this region. Furthermore, Table S6 summarizes the mutated regions associated with other signatures active in CLL. SBS1, SBS5, and SBS40 were all considered clock-like signatures [25]; therefore, it was unlikely that these signatures would act on specific regions of the genome. As shown in Table S6, the number of mutated regions associated with these signatures was reasonably small compared to that for SBS9, the mutational process for which a specific target region existed.

**Table 3:** Frequently mutated regions in clones with high SBS9 activity for CLL samples.

| Hugo symbol | Variant classification | p-value < FWER |
| --- | --- | --- |
| IGHJ6 | RNA | 1.22E-33 |
| IGLL5 | 5'Flank | 8.13E-18 |
| IGLL5 | Intron | 5.82E-17 |
| BCL6 | Intron | 4.25E-15 |
| AC018717.1 | lincRNA | 3.98E-14 |
| AC096579.7 | RNA | 3.60E-11 |
| IGLV3-25 | RNA | 3.82E-11 |
| IGLV3-1 | RNA | 1.08E-10 |
| IGKJ2 | RNA | 5.53E-10 |
| KIAA0125 | 5'Flank | 8.89E-10 |
| Unknown | IGR | 1.05E-09 |
| IGKV4-1 | RNA | 1.76E-09 |
| DMD | Intron | 1.24E-08 |
| TCL1A | Intron | 1.60E-08 |
| IMMP2L | Intron | 6.96E-08 |
| FSTL5 | Intron | 1.12E-07 |
| IGKC | RNA | 2.72E-07 |
| IGKV1-5 | RNA | 2.81E-07 |
| IGKJ3 | RNA | 4.28E-07 |
| BCL2 | 5'UTR | 1.42E-06 |
| IGHD3-10 | RNA | 1.58E-06 |
| FHIT | Intron | 1.73E-06 |

This table lists the mutated regions significantly correlated with SBS9 in CLL samples detected according to the test pipeline. The left column indicates the gene name, and the middle column the region of the gene. They are listed in order of decreasing p-value, and only those that were determined to be significant by Bonferroni correction are shown.

We also applied the same test to BNHL results to investigate the relationship between signatures and mutations in blood cancer. Table S7 shows the mutated regions that were significantly enriched with clones with high SBS9 activity. Combined with Table 3, the FWER-based test yielded only one region associated with SBS9 in common with CLL, which was an intronic region of FHIT. In addition, the mutations in the immunoglobulin-coding region were not enriched in the clone with high SBS9 activity for BNHL samples, which significantly differed from the result obtained for CLL samples.

Then, we focused on the mutated regions in both CLL and BNHL samples among those detected using a simple significance level *α* = 0.05 rather than FWER. This approach did not include multiple testing corrections, which might have led to false positives. However, this ensured a certain degree of reliability because two results that were obtained independently were compared. All four signatures (SBS1, SBS5, SBS9, and SBS40) active in the CLL samples were also active in some of the BNHL samples, and we focused on these signatures. We have summarized the number of significantly mutated regions and their Hugo symbols in Table S8 and S9. The number of mutated regions associated with SBS9 was high compared to other signatures, and interestingly, a missense mutation for KLHL6 was identified in the gene set corresponding to SBS9; thus, we could hypothesize that the etiology of SBS9, probably AID or polymerase eta, might be associated with KLHL6. Recent studies suggest that KLHL6 may be an essential tumor suppressor gene in B-cell lymphoma [26, 27]. Therefore, KLHL6 aberration may lead to high activity of the mutational process of SBS9, resulting in a large number of somatic mutations. However, further validations are required to reveal if this hypothesis is true because we only considered somatic mutations not including germline mutations lying on the actual genome in this study.

We performed gene ontology (GO) analysis on the resulting gene set to determine if there were any common features of each signature (GO Enrichment Analysis powered by PANTHER: http://geneontology.org/). Thus, significant terms were detected only for the gene set corresponding to SBS9, and the most interesting ones are shown in Table 4. For example, “cell junction organization” and “cell adhesion” annotations were detected, and they included cadherin-coding regions. These functions are closely related to the immune system, and cell adhesion is related to whether or not a tumor can acquire metastatic potential [28, 29]; hence, SBS9 activity might also be closely associated with a poor prognosis of blood cancer. Notably, these mutations also occurred in BNHL samples, compared to the coding region of immunoglobulin, where mutations were concentrated only in CLL samples. Because mutations on these genes were concentrated in the intronic region, we suspected that they were the consequence of SBS9 rather than the cause. Additionally, Figure S3 shows a Venn diagram of the overlaps between signatures, indicating that the mutated regions for SBS9 were particularly unique.

**Table 4:**
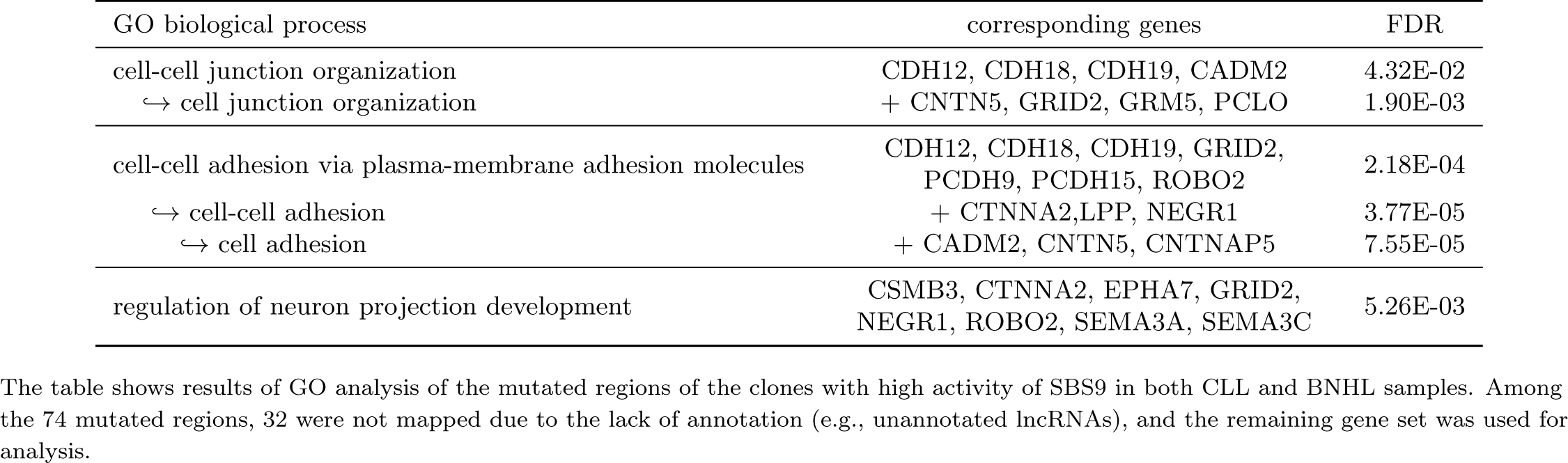
Significant GO terms related with SBS9 in both CLL and BNHL samples.

### Future work

SigTracer, which performs clone decomposition based on mutation signatures, showed significant potential of providing novel insights into mutational processes because of its improved accuracy of assigning mutations to signatures by considering VAF. However, the method suffers from some limitations. An issue in the modeling is that there is no established method that can predict the correct number of clones for arbitrary data. In our simulation experiments, we showed that the model achieved a sufficiently high likelihood with fewer clones than the true number for low-coverage data, which highlighted the limitation in terms of the amount of information in the input. To solve this problem, it is necessary to incorporate other data sources that are effective for inference in addition to mutation types and VAF.

Moreover, there is room for reconsideration of the type of signature used in the analysis of real datasets. SBS84 and SBS85, the existence of which has been demonstrated in the latest studies [21], are signatures known to be active in blood cancer. The inclusion of these signatures may lead to novel findings. When more signatures are included as active signature candidates, a phenomenon called “signature bleeding” occurs, in which signatures that are not actually active are erroneously presumed to be active [30]. To avoid such a situation, we must introduce a regularization to the model to reduce the number of active signatures used in the fitting method, such as sigLASSO [18].

Regarding the current test pipeline, we can detect the causality of mutations due to signatures with no difficulty using this method; however, causal inference to signatures due to mutations (i.e., the identification of the driver mutations) needs to be performed by considering the evolutionary background of the tumor. For example, if tumors follow a branching evolutionary process [31], each tumor cell may have mutations that originate from multiple clones. In such cases, it is difficult to capture the causality of signatures due to mutations because there is a possibility that the active signature in a certain clone is activated by mutations belonging to other clones. Thus, the causality of signatures due to mutations can be detected using the method described here only if the tumor evolution follows a specific process, such as the Big Bang dynamics which state that cancer cells evolve independently [32], where each clone directly represents a mutational population carried by each cell. Recent reports concerning colorectal cancer support the neutral evolution and the Big Bang model [33, 34], and future accumulation of knowledge on cancer evolution will resolve this issue.

## CONCLUSION

We developed SigTracer, a signature-based method for estimating clonal evolution based on mutation types and VAF observed via bulk sequencing. In computational simulations, SigTracer outperformed the existing method in terms of model selection and accuracy for ideal artificial data. In addition, we applied SigTracer to CLL samples; the results were consistent with previous findings that SBS9, which is associated with AID or polymerase eta, intensively mutates the immunoglobulin-coding regions. Furthermore, we performed the same analysis on BNHL samples using SigTracer and found that SBS9 also includes intensive mutation of the regions coding for cadherins and other genes regulating cell adhesion. Our results indicate that AID or polymerase eta activity may be induced in more regions related to the immune system than previously known. These new observations were obtained because of the improved accuracy of assigning mutations to signatures by considering not only mutation types but also VAF. We believe applying the proposed method to other cancer types may lead to the annotation of signatures for which mutational processes and target regions are unknown. Our results provide an excellent prospect for understanding the mechanism of carcinogenesis.

## DATA AVAILABILITY

Our implementation of SigTracer in C++ and custom scripts in Python is available at GitHub repository: https://github.com/qkirikigaku/SigTracer.

## ACKNOWLEDGMENTS

Computation for this study was partially performed on the NIG supercomputer at ROIS National Institute of Genetics. We thank members at Hamada Laboratory for the valuable discussions regarding this study.

## FUNDING

This work was supported by the Ministry of Education, Culture, Sports, Science and Technology (KAKENHI) [grant numbers JP17K20032, JP16H05879, JP16H01318, JP16H02484,JP16H06279 and JP20H00624 to MH, JP20J20016 to TM].

## Conflict of interest statement

None declared.

## Author contributions

TM conceived this study. MH supervised this study. TM designed the algorithms, implemented the method, and performed all the computational experiments. TM and MH analyzed the results. TM wrote the draft. MH revised the manuscript critically. All authors read and approved the final manuscript.

## Notes

https://github.com/qkirikigaku/SigTracer

### Competing Interest Statement

The authors have declared no competing interest.

