## Supplementary Materials for "Clone decomposition based on mutation signatures provides novel insights into mutational processes"

**Taro Matsutani**[1,2,*] **and Michiaki Hamada**[1,2,3,*]

[1]Graduate School of Advanced Science and Engineering, Waseda University, 55N-06-10, 3-4-1, Okubo Shinjuku-ku, Tokyo 169–8555, Japan
[2]Computational Bio Big-Data Open Innovation Laboratory (CBBD-OIL), National Institute of Advanced Industrial Science and Technology (AIST), Tokyo 169–8555, Japan
[3]Graduate School of Medicine, Nippon Medical School, Sendagi, Bunkyo, Tokyo 113-8602, Japan

#### 1 Derivation of the algorithm to estimate parameters in SigTracer

**Table S1.** Notations utilized in the SigTracer model.

| Constants / Observed variables | Definitions |
|---|---|
| $N$ | The number of mutations in the sample |
| $J$ | The number of clones in the sample |
| $V$ | The number of mutation types ($V = 96$ in this study; considering tri-nucleotide) |
| $K$ | The number of mutation signatures appeared in the sample |
| $x_n$ | The mutation type (represented in trinucleotide) for the $n$-th mutation $(1 \leq n \leq N)$ |
| $B_n$ | The sequenced number of the reads with mutated allele for the locus of the $n$-th mutation |
| $D_n$ | The sequenced number of total reads for the locus of the $n$-th mutation |
| $\boldsymbol{\phi}_k = \{\phi_{k,v}\}_{v=1}^{V}$ | The mutational distribution for $k$-th signature $(1 \leq k \leq K)$ |
| $P$ | The proportion of the reads from cancer cell in all the sequenced reads (i.e. purity) |
| $C_n^{(\text{tumor})}$ | The copy number of cancer cell for the locus of the $n$-th mutation |
| $C_n^{(\text{major})}$ | The copy number of major allele in cancer cell for the locus of the $n$-th mutation |
| $C_n^{(\text{normal})}$ | The copy number of normal cell for the locus of the $n$-th mutation |
| **Hidden variables** | **Definitions** |
| $z_n$ | Index to show the signature that generates the $n$-th mutation $(1 \leq z_n \leq K)$ |
| $U_n$ | Index to show the clone carrying the $n$-th mutation $(1 \leq U_n \leq J)$ |
| $M_n$ | The copy number of mutated allele in cancer cell for the locus of the $n$-th mutation $(1 \leq M_n \leq C_n^{(\text{major})})$ |
| **Parameters** | **Definitions** |
| $\lambda_j$ | The cancer cell fraction (CCF) for the $j$-th clone $(1 \leq j \leq J)$ |
| $\rho_j$ | The overdispersion parameter for the $j$-th clone |
| $\tau$ | The hyperparameter of $\boldsymbol{\rho} = \{\rho_j\}_{j=1}^{J}$ |
| $\pi_j$ | The proportion of the mutations belonging to the $j$-th clone in all the mutations |
| $\boldsymbol{\alpha}^{(\pi)} = \{\alpha_j^{(\pi)}\}_{j=1}^{J}$ | The hyperparameters to control $\boldsymbol{\pi} = \{\pi_j\}_{j=1}^{J}$ |
| $\boldsymbol{\theta}_j = \{\theta_{j,k}\}_{k=1}^{K}$ | The signature activity for the $j$-th clone |
| $\boldsymbol{\alpha}_j^{(\theta)} = \{\alpha_{j,k}^{(\theta)}\}_{k=1}^{K}$ | The hyperparameters to control $\boldsymbol{\theta}_j$ |

In this section, we provide a detailed derivation of the update formula that appears in the algorithm for estimating the parameters of SigTracer introduced in the main text.

##### 1.1 CVB inference

As described in the main text, ELBO, the objective function in collapsed CVB, can be defined as follows:

$$F[q(\boldsymbol{z},\boldsymbol{U},\boldsymbol{M})] \equiv \sum_{n=1}^{N}\sum_{z_n}\sum_{U_n}\sum_{M_n} q(z_n,U_n,M_n)\log\frac{p(x_n,B_n,z_n,U_n,M_n \mid D_n,C_n,P,\boldsymbol{\lambda},\boldsymbol{\rho},\boldsymbol{\alpha}^{(\pi)},\boldsymbol{\alpha}^{(\theta)},\boldsymbol{\phi})}{q(z_n,U_n,M_n)}.$$

It is impossible to compute this value explicitly; however, by focusing only on the terms involved in $q(z_n,U_n,M_n)$ (let it $\tilde{F}[q(z_n,U_n,M_n)]$), we can derive the update formula. Assuming that all mutations are independent, $\tilde{F}[q(z_n,U_n,M_n)]$ is calculated as follows:

$$\begin{aligned}\tilde{F}[q(z_n,U_n,M_n)] = &\sum_{n=1}^{N}\sum_{z_n}\sum_{U_n}\sum_{M_n} q(z_n,U_n,M_n)q(\boldsymbol{z}^{\setminus n},\boldsymbol{U}^{\setminus n},\boldsymbol{M}^{\setminus n})\\ &\times\log\frac{p(x_n,B_n,z_n,U_n,M_n \mid \boldsymbol{z}^{\setminus n},\boldsymbol{U}^{\setminus n},\boldsymbol{D}_n,C_n,P,\boldsymbol{\lambda},\boldsymbol{\rho},\boldsymbol{\alpha}^{(\pi)},\boldsymbol{\alpha}^{(\theta)},\boldsymbol{\phi})}{q(z_n,U_n,M_n)},\end{aligned} \tag{S1}$$

where the variables with $\setminus n$ show the parameters excluding the $n$-th value. In Eq. (S1), the joint probability for mutation $n$ can be obtained as follows:

$$\begin{aligned}&\log p(x_n,B_n,z_n=k,U_n=j,M_n=m \mid \boldsymbol{z}^{\setminus n},\boldsymbol{U}^{\setminus n},D_n,C_n,P,\boldsymbol{\lambda},\boldsymbol{\rho},\boldsymbol{\alpha}^{(\pi)},\boldsymbol{\alpha}^{(\theta)},\boldsymbol{\phi})\\ &\propto \log p(x_n \mid \phi_k)+\log p(B_n \mid, U_n=j,M_n=m,D_n,C_n,P,\lambda_j,\rho_j)+\log p(z_n=k \mid U_n=j,\boldsymbol{z}^{\setminus n},\boldsymbol{U}^{\setminus n},\boldsymbol{\alpha}^{(\theta)})+\log p(U_n=j \mid \boldsymbol{U}^{\setminus n},\boldsymbol{\alpha}^{(\pi)})\\ &=\log\phi_{k,x_n}+\log\frac{\Gamma(B_n+\mu_{n,j,m})\Gamma(D_n-B_n+\nu_{n,j,m})\Gamma(\mu_{n,j,m}+\nu_{n,j,m})}{\Gamma(D_n+\mu_{n,j,m}+\nu_{n,j,m})\Gamma(\nu_{n,j,m})\Gamma(\mu_{n,j,m})}+\log\frac{\alpha_{j,k}^{(\theta)}+\kappa_{j,k}^{\setminus n}}{\sum_{k'}\left[\alpha_{j,k'}^{(\theta)}+\kappa_{j,k'}^{\setminus n}\right]}+\log\frac{\alpha_{j}^{(\pi)}+\kappa_{j}^{\setminus n}}{\sum_{j'}\left[\alpha_{j'}^{(\pi)}+\kappa_{j'}^{\setminus n}\right]},\end{aligned} \tag{S2}$$

where $\mu_{n,j,m}$ shows $\rho_j\lambda_j\eta_{n,m}$ and $\nu_{n,j,m}$ shows $\rho_j(1-\lambda_j\eta_{n,m})$, respectively (see Eq. (1) in the main text for the definition of $\eta_{n,m}$). In addition, $\kappa_j$ shows the number of mutations that belong to the $j$-th clone, and $\kappa_{j,k}$ is the number of mutations that belong to the $j$-th clone and are caused by $k$-th signature. We substitute Eq. (S2) into Eq. (S1) and differentiate by $q(z_n=k,U_n=j,M_n=m)$ to obtain the stationary point:

$$\begin{aligned}q(z_n=k,U_n=j,M_n=m) \propto \exp\Bigg[&\log\phi_{k,x_n}+\log\frac{\Gamma(B_n+\mu_{n,j,m})\Gamma(D_n-B_n+\nu_{n,j,m})\Gamma(\mu_{n,j,m}+\nu_{n,j,m})}{\Gamma(D_n+\mu_{n,j,m}+\nu_{n,j,m})\Gamma(\nu_{n,j,m})\Gamma(\mu_{n,j,m})}\\ &+\mathbb{E}_{q(\boldsymbol{z}^{\setminus n},\boldsymbol{U}^{\setminus n},\boldsymbol{M}^{\setminus n})}\left\{\log\frac{\alpha_{j,k}^{(\theta)}+\kappa_{j,k}^{\setminus n}}{\sum_{k'}\left\{\alpha_{j,k'}^{(\theta)}+\kappa_{j,k'}^{\setminus n}\right\}}\right\}+\mathbb{E}_{q(\boldsymbol{z}^{\setminus n},\boldsymbol{U}^{\setminus n},\boldsymbol{M}^{\setminus n})}\left\{\log\frac{\alpha_{j}^{(\pi)}+\kappa_{j}^{\setminus n}}{\sum_{j'}\left\{\alpha_{j'}^{(\pi)}+\kappa_{j'}^{\setminus n}\right\}}\right\}\Bigg]\\ =\exp\Bigg[&\log\phi_{k,x_n}+\log\frac{\Gamma(B_n+\mu_{n,j,m})\Gamma(D_n-B_n+\nu_{n,j,m})\Gamma(\rho_j)}{\Gamma(D_n+\rho_j)\Gamma(\nu_{n,j,m})\Gamma(\mu_{n,j,m})}\\ &+\log\frac{\alpha_{j,k}^{(\theta)}+\sum_{n'\neq n}\sum_{M_{n'}}q(z_{n'}=k,U_{n'}=j,M_{n'})}{\sum_{k'}\left\{\alpha_{j,k'}^{(\theta)}+\sum_{n'\neq n}\sum_{M_{n'}}q(z_{n'}=k',U_{n'}=j,M_{n'})\right\}}\\ &+\log\frac{\alpha_{j}^{(\pi)}+\sum_{n'\neq n}\sum_{z_{n'}}\sum_{M_{n'}}q(z_{n'},U_{n'}=j,M_{n'})}{\sum_{j'}\left\{\alpha_{j'}^{(\pi)}+\sum_{n'\neq n}\sum_{z_{n'}}\sum_{M_{n'}}q(z_{n'},U_{n'}=j',M_{n'})\right\}}\Bigg].\end{aligned} \tag{S3}$$

Note that the zero-order Taylor expansion of $\mathbb{E}[\log x]=\log\mathbb{E}[x]$ is used to calculate the expected value against $q(\boldsymbol{z}^{\setminus n},\boldsymbol{U}^{\setminus n},\boldsymbol{M}^{\setminus n})$. Finally, by normalizing with respect to $z_n,U_n,M_n$, we can obtain the responsibility, $q(z_n=k,U_n=j,M_n=m)$.

#### 1.2 Variational Bayes method for model selection

To select the number of clones, $J$, SigTracer first infers the parameters with a VB and uses the resulting ELBO value. In ELBO with VB, the approximate posteriors, $q(\boldsymbol{\pi})$ and $q(\boldsymbol{\theta})$, were not marginalized and ELBO was defined as follows:

$$
\begin{aligned}
F_{\mathrm{VB}}[q(\boldsymbol{z},\boldsymbol{U},\boldsymbol{M},\boldsymbol{\pi},\boldsymbol{\theta})] &\equiv \int_{\boldsymbol{\pi}}\int_{\boldsymbol{\theta}} q(\boldsymbol{\pi})q(\boldsymbol{\theta}) \sum_{n=1}^{N}\sum_{z_n}\sum_{U_n}\sum_{M_n} q(z_n,U_n,M_n)\log\frac{p(x_n,B_n,z_n,U_n,M_n,\boldsymbol{\pi},\boldsymbol{\theta}\mid D_n,C_n,P,\boldsymbol{\lambda},\boldsymbol{\rho},\boldsymbol{\alpha}^{(\pi)},\boldsymbol{\alpha}^{(\theta)},\boldsymbol{\phi})}{q(z_n,U_n,M_n)} d\boldsymbol{\pi} d\boldsymbol{\theta} \\
&= \sum_{n=1}^{N}\sum_{z_n}\sum_{U_n}\sum_{M_n} q(z_n=k,U_n=j,M_n=m)\log p(x_n\mid\boldsymbol{\phi}_k) \\
&+ \sum_{n=1}^{N}\sum_{z_n}\sum_{U_n}\sum_{M_n} q(z_n=k,U_n=j,M_n=m)\log p(B_n\mid U_n=j,M_n=m,D_n,C_n,\rho_j,\lambda_j) \\
&+ \sum_{n=1}^{N}\sum_{z_n}\sum_{U_n}\sum_{M_n} q(z_n=k,U_n=j,M_n=m)\int q(\boldsymbol{\theta}_j)\log p(z_n=k\mid\boldsymbol{\theta}_j)d\boldsymbol{\theta}_j \\
&+ \sum_{n=1}^{N}\sum_{z_n}\sum_{U_n}\sum_{M_n} q(z_n=k,U_n=j,M_n=m)\int q(\boldsymbol{\pi})\log p(U_n=j\mid\boldsymbol{\pi})d\boldsymbol{\pi} \\
&+ \sum_{j=1}^{J}\int q(\boldsymbol{\theta}_j)\log\frac{p(\boldsymbol{\theta}_j\mid\boldsymbol{\alpha}^{(\theta)})}{q(\boldsymbol{\theta}_j)}d\boldsymbol{\theta}_j + \int q(\boldsymbol{\pi})\log\frac{p(\boldsymbol{\pi}\mid\boldsymbol{\alpha}^{(\pi)})}{q(\boldsymbol{\pi})}d\boldsymbol{\pi} \\
&- \sum_{n=1}^{N}\sum_{z_n}\sum_{U_n}\sum_{M_n} q(z_n=k,U_n=j,M_n=m)\log q(z_n=k,U_n=j,M_n=m). \qquad \text{(S4)}
\end{aligned}
$$

The integrals for $\boldsymbol{\pi}$ and $\boldsymbol{\theta}$ in $F_{\mathrm{VB}}[q(\boldsymbol{z},\boldsymbol{U},\boldsymbol{M},\boldsymbol{\pi},\boldsymbol{\theta})]$ can be developed from the conjugacy between the Dirichlet and categorical distributions as follows:

$$
\begin{aligned}
&\int q(\boldsymbol{\theta}_j)\log p(z_n=k\mid\boldsymbol{\theta}_j)d\boldsymbol{\theta}_j = \Psi(\xi^{\theta}_{j,k}) - \Psi(\sum_{k'=1}^{K}\xi^{\theta}_{j,k'}) \ \text{ with } \ \xi^{\theta}_{j,k} = \sum_{n=1}^{N}\sum_{m=1}^{C_n^{(\mathrm{major})}} q(z_n=k,U_n=j,M_n=m) + \alpha^{(\theta)}_{j,k}, \\
&\int q(\boldsymbol{\pi})\log p(U_n=j\mid\boldsymbol{\pi})d\boldsymbol{\pi} = \Psi(\xi^{\pi}_{j}) - \Psi(\sum_{j'=1}^{J}\xi^{\pi}_{j'}) \ \text{ with } \ \xi^{\pi}_{j} = \sum_{n=1}^{N}\sum_{k=1}^{K}\sum_{m=1}^{C_n^{(\mathrm{major})}} q(z_n=k,U_n=j,M_n=m) + \alpha^{(\pi)}_{j},
\end{aligned}
$$

where $\Psi(\cdot)$ shows the digamma function. Similar to CVB, we extract the terms of $F_{\mathrm{VB}}[q(\boldsymbol{z},\boldsymbol{U},\boldsymbol{M},\boldsymbol{\pi},\boldsymbol{\theta})]$ involved in $q(z_n,U_n,M_n)$:

$$
\begin{aligned}
&\tilde{F}_{\mathrm{VB}}[q(z_n=k,U_n=j,M_n=m)] \\
&= q(z_n=k,U_n=j,M_n=m)\Bigg[\log p(x_n\mid\boldsymbol{\phi}_k) + \log p(B_n\mid U_n=j,M_n=m,D_n,C_n,\rho_j,\lambda_j) + \left\{\Psi(\xi^{\theta}_{j,k}) - \Psi(\sum_{k'=1}^{K}\xi^{\theta}_{j,k'})\right\} \\
&+ \left\{\Psi(\xi^{\pi}_{j}) - \Psi(\sum_{j'=1}^{J}\xi^{\pi}_{j'})\right\} - \log q(z_n=k,U_n=j,M_n=m)\Bigg].
\end{aligned}
$$

Then, we take the stationary point:

$$
\begin{aligned}
q(z_n=k,U_n=j,M_n=m) \propto \exp\Bigg[&\log\phi_{k,x_n} + \log\frac{\Gamma(B_n+\mu_{n,j,m})\Gamma(D_n-B_n+\nu_{n,j,m})\Gamma(\mu_{n,j,m}+\nu_{n,j,m})}{\Gamma(D_n+\mu_{n,j,m}+\nu_{n,j,m})\Gamma(\nu_{n,j,m})\Gamma(\mu_{n,j,m})} \\
&+ \left\{\Psi(\xi^{\theta}_{j,k}) - \Psi(\sum_{k'=1}^{K}\xi^{\theta}_{j,k'})\right\} + \left\{\Psi(\xi^{\pi}_{j}) - \Psi(\sum_{j'=1}^{J}\xi^{\pi}_{j'})\right\}\Bigg]. \qquad \text{(S5)}
\end{aligned}
$$

For $q(\boldsymbol{\theta}_j)$, ELBO is transformed as follows:

$$
\tilde{F}_{\mathrm{VB}}[q(\boldsymbol{\theta}_j)] = \int q(\boldsymbol{\theta}_j)\left\{\sum_{n=1}^{N}\sum_{k=1}^{K}\sum_{m=1}^{C_n^{(\mathrm{major})}} q(z_n=k,U_n=j,M_n=m)\log p(z_n\mid\boldsymbol{\theta}_j) + \log\frac{p(\boldsymbol{\theta}_j\mid\boldsymbol{\alpha}^{(\theta)})}{q(\boldsymbol{\theta}_j)}\right\}d\boldsymbol{\theta}_j.
$$

Denote $\tilde{F}_{\mathrm{VB}}[q(\boldsymbol{\theta}_j)] = \int q(\boldsymbol{\theta}_j) f(q(\boldsymbol{\theta}_j)) d\boldsymbol{\theta}_j$ for the above equation; then, $q(\boldsymbol{\theta}_j)$ satisfying $\frac{\partial f(q(\boldsymbol{\theta}_j))}{q(\boldsymbol{\theta}_j)} = 0$ automatically satisfies $\frac{\partial \tilde{F}_{\mathrm{VB}}(q(\boldsymbol{\theta}_j))}{\partial q(\boldsymbol{\theta}_j)} = 0$. Thus, we solve for $\frac{\partial f(q(\boldsymbol{\theta}_j))}{\partial q(\boldsymbol{\theta}_j)} = 0$ and get

$$q(\boldsymbol{\theta}_j) \propto p(\boldsymbol{\theta}_j \mid \boldsymbol{\alpha}^{(\theta)}) \exp\left\{ \sum_{n=1}^{N} \sum_{k=1}^{K} \sum_{m=1}^{C_n^{(\mathrm{major})}} q(z_n = k, U_n = j, M_n = m) \log p(z_n \mid \boldsymbol{\theta}_j) \right\} \propto \prod_{k=1}^{K} \theta_{j,k}^{\xi_{j,k}^{\theta} - 1}. \tag{S6}$$

Thus, the approximate posterior distribution becomes a functional form of the Dirichlet distribution, and its parameter is $\xi_{j,k}^{\theta}$. For $q(\boldsymbol{\pi})$, in the same way as for $q(\boldsymbol{\theta})$, we can obtain following update formula:

$$q(\boldsymbol{\pi}) \propto \prod_{j=1}^{J} \pi_j^{\xi_j^{\pi} - 1}. \tag{S7}$$

By substituting all obtained estimators into Eq. (S4), we can obtain the value which is the criterion to select the number of clones, $J$.

##### 1.3 Fixed-point iteration to estimate hyper-parameters

We predicted the hyper-parameters including $\boldsymbol{\alpha}^{(\pi)}$, $\boldsymbol{\alpha}^{(\theta)}$, $\boldsymbol{\lambda}$, and $\boldsymbol{\rho}$ using the fixed-point iteration method. In this method, we first derived the lower bound of ELBO using the gamma function and computed the stationary points for each parameter.

First, for $\boldsymbol{\alpha}^{(\pi)}$, the ELBO terms involved in $\boldsymbol{\alpha}^{(\pi)}$ were extracted as follows:

$$\tilde{F}[\boldsymbol{\alpha}^{(\pi)}] = \log \frac{\Gamma(\sum_{j=1}^{J} \alpha_j^{(\pi)})}{\prod_{j=1}^{J} \Gamma(\alpha_j^{(\pi)})} - \log \frac{\Gamma(N + \sum_{j=1}^{J} \alpha_j^{(\pi)})}{\prod_{j=1}^{J} \Gamma(\kappa_j + \alpha_j^{(\pi)})} = \log \frac{\Gamma(\sum_{j=1}^{J} \alpha_j^{(\pi)})}{\Gamma(N + \sum_{j=1}^{J} \alpha_j^{(\pi)})} + \sum_{j=1}^{J} \log \frac{\Gamma(\kappa_j + \alpha_j^{(\pi)})}{\Gamma(\alpha_j^{(\pi)})},$$

and their lower bound was calculated with $\hat{\alpha}_j^{(\pi)}$ denoted as $\alpha_j^{(\pi)}$ before the update, as shown below:

$$\tilde{F}[\alpha_j^{(\pi)}] \geq -\alpha_j^{(\pi)} \left\{ \Psi\left(N + \sum_{j=1}^{J} \hat{\alpha}_j^{(\pi)}\right) - \Psi\left(\sum_{j=1}^{J} \hat{\alpha}_j^{(\pi)}\right) \right\} + \log \alpha_j^{(\pi)} [\hat{\alpha}_j^{(\pi)} \{\Psi(\kappa_j + \hat{\alpha}_j^{(\pi)}) - \Psi(\hat{\alpha}_j^{(\pi)})\}] + (\text{const.}).$$

Then, $\alpha_j^{(\pi)}$ could be updated as follows:

$$\begin{aligned} &\frac{\hat{\alpha}_j^{(\pi)} \{\Psi(\kappa_j + \hat{\alpha}_j^{(\pi)}) - \Psi(\hat{\alpha}_j^{(\pi)})\}}{\alpha_j^{(\pi)}} - \left\{ \Psi\left(N + \sum_{j=1}^{J} \hat{\alpha}_j^{(\pi)}\right) - \Psi\left(\sum_{j=1}^{J} \hat{\alpha}_j^{(\pi)}\right) \right\} = 0 \\ \Leftrightarrow &\alpha_j^{(\pi)} = \frac{\hat{\alpha}_j^{(\pi)} \{\Psi(\kappa_j + \hat{\alpha}_j^{(\pi)}) - \Psi(\hat{\alpha}_j^{(\pi)})\}}{\Psi\left(N + \sum_{j=1}^{J} \hat{\alpha}_j^{(\pi)}\right) - \Psi\left(\sum_{j=1}^{J} \hat{\alpha}_j^{(\pi)}\right)}. \end{aligned} \tag{S8}$$

We obtained the updated formula for $\boldsymbol{\alpha}^{(\theta)}$ in the same manner for $\boldsymbol{\alpha}^{(\pi)}$ with $\hat{\alpha}_{j,k}^{(\theta)}$ denoted as $\alpha_{j,k}^{(\theta)}$ before the update, as follows:

$$\alpha_{j,k}^{(\theta)} = \frac{\hat{\alpha}_{j,k}^{(\theta)} \sum_{j=1}^{J} \sum_{k=1}^{K} [\Psi(\kappa_{j,k} + \hat{\alpha}_{,j,k}^{(\theta)}) - \Psi(\hat{\alpha}_{,j,k}^{(\theta)})]}{K \sum_{j=1}^{J} [\Psi(\kappa_j + K\hat{\alpha}_{j,k}^{(\theta)}) - \Psi(K\hat{\alpha}_{j,k}^{(\theta)})]}. \tag{S9}$$

Next, we introduced how to estimate $\boldsymbol{\lambda}$ and $\boldsymbol{\rho}$ in BetaBinomial distribution with fixed-point iterations. The ELBO terms related to $\lambda_j$ were as follows:

$$\tilde{F}[\lambda_j] = \sum_{n=1}^{N} \sum_{k=1}^{K} \sum_{m=1}^{C_n^{(\mathrm{major})}} q(z_n = k, U_n = j, M_n = m) \log \frac{\Gamma(B_n + \mu_{n,j,m}) \Gamma(D_n - B_n + \nu_{n,j,m})}{\Gamma(\mu_{n,j,m}) \Gamma(\nu_{n,j,m})}.$$

We then take the lower bound of $\log \Gamma(B_n + \mu_{n,j,m}) - \log \Gamma(\mu_{n,j,m})$ and $\log \Gamma(D_n - B_n + \nu_{n,j,m}) - \log \Gamma(\nu_{n,j,m})$ with $\hat{\lambda}_j$, which is denoted as $\lambda_j$ before update:

$$\begin{aligned} \log \frac{\Gamma(B_n + \mu_{n,j,m})}{\log \Gamma(\mu_{n,j,m})} &\to \hat{\mu}_{n,j,m}^{\lambda} \{\Psi(B_n + \hat{\mu}_{n,j,m}^{\lambda}) - \Psi(\hat{\mu}_{n,j,m}^{\lambda})\} \log \lambda_j + (\text{const.}), \\ \log \frac{\Gamma(D_n - B_n + \nu_{n,j,m})}{\log \Gamma(\nu_{n,j,m})} &\to \hat{\nu}_{n,j,m}^{\lambda} \{\Psi(D_n - B_n + \hat{\nu}_{n,j,m}^{\lambda}) - \Psi(\hat{\nu}_{n,j,m}^{\lambda})\} \log(1 - \lambda_j \eta_{n,m}) + (\text{const.}), \end{aligned}$$

where

$$\hat{\mu}^{\lambda}_{n,j,m} = \rho_j \hat{\lambda}_j \eta_{n,m},$$
$$\hat{\nu}^{\lambda}_{n,j,m} = \rho_j (1 - \hat{\lambda}_j \eta_{n,m}).$$

Substituting them into ELBO and formulating the equation for $\partial \tilde{F}[\lambda_j]/\partial \lambda_j = 0$:

$$\sum_{n=1}^{N} \sum_{k=1}^{K} \sum_{m=1}^{C_n^{(\text{major})}} q(z_n = k, U_n = j, M_n = m)$$
$$\times \left[ \frac{\hat{\mu}^{\lambda}_{n,j,m} \{\Psi(B_n + \hat{\mu}^{\lambda}_{n,j,m}) - \Psi(\hat{\mu}^{\lambda}_{n,j,m})\}}{\lambda_j} - \frac{\eta_{n,m} \hat{\nu}^{\lambda}_{n,j,m} \{\Psi(D_n - B_n + \hat{\nu}^{\lambda}_{n,j,m}) - \Psi(\hat{\nu}^{\lambda}_{n,j,m})\}}{1 - \lambda_j \eta_{n,m}} \right] = 0$$
$$\Leftrightarrow \lambda_j = \frac{\sum_{n=1}^{N} \sum_{k=1}^{K} \sum_{m=1}^{C_n^{(\text{major})}} q(z_n = k, U_n = j, M_n = m) \Xi^{\lambda}_{n,j,m}}{\sum_{n=1}^{N} \sum_{k=1}^{K} \sum_{m=1}^{C_n^{(\text{major})}} q(z_n = k, U_n = j, M_n = m) \eta_{n,m} (\Xi^{\lambda}_{n,j,m} + \Phi^{\lambda}_{n,j,m})}, \tag{S10}$$

where

$$\Xi^{\lambda}_{n,j,m} = \hat{\mu}^{\lambda}_{n,j,m} \{\Psi(B_n + \hat{\mu}^{\lambda}_{n,j,m}) - \Psi(\hat{\mu}^{\lambda}_{n,j,m})\},$$
$$\Phi^{\lambda}_{n,j,m} = \hat{\nu}^{\lambda}_{n,j,m} \{\Psi(D_n - B_n + \hat{\nu}^{\lambda}_{n,j,m}) - \Psi(\hat{\nu}^{\lambda}_{n,j,m})\}.$$

Finally, the ELBO terms related to $\rho_j$ are given as follows:

$$\tilde{F}[\rho_j] = \sum_{n=1}^{N} \sum_{k=1}^{K} \sum_{m=1}^{C_n^{(\text{major})}} q(z_n = k, U_n = j, M_n = m) \log \frac{\Gamma(B_n + \mu_{n,j,m}) \Gamma(D_n - B_n + \nu_{n,j,m}) \Gamma(\rho_j)}{\Gamma(\mu_{n,j,m}) \Gamma(\nu_{n,j,m}) \Gamma(D_n + \rho_j)}.$$

In the update of $\rho_j$, we set the exponential distribution $p(\rho \mid \tau)$ as the prior distribution to suppress overfitting:

$$\log p(\rho \mid \tau) = \log \tau - \tau \rho.$$

In addition, the lower bound of the terms related with $\rho_j$ was calculated as follows:

$$\log \frac{\Gamma(B_n + \mu^{\rho}_{n,j,m})}{\log \Gamma(\mu^{\rho}_{n,j,m})} \to \log \rho_j \{\Psi(B_n + \hat{\mu}^{\rho}_{n,j,m}) - \Psi(\hat{\mu}^{\rho}_{n,j,m})\} + (\text{const.}),$$
$$\log \frac{\Gamma(D_n - B_n + \nu^{\rho}_{n,j,m})}{\log \Gamma(\nu^{\rho}_{n,j,m})} \to \log \rho_j \{\Psi(D_n - B_n + \hat{\nu}^{\rho}_{n,j,m}) - \Psi(\hat{\nu}^{\rho}_{n,j,m})\} + (\text{const.}),$$
$$\log \frac{\Gamma(\rho_j)}{\Gamma(D_n + \rho_j)} \to -\rho_j \{\Psi(D_n + \hat{\rho}_j) - \Psi(\hat{\rho}_j)\} + (\text{const.}),$$

where

$$\hat{\mu}^{\rho}_{n,j,m} = \hat{\rho}_j \lambda_j \eta_{n,m},$$
$$\hat{\nu}^{\rho}_{n,j,m} = \hat{\rho}_j (1 - \lambda_j \eta_{n,m}).$$

Combining them , the objective function is given as follows:

$$\tilde{F}[\rho_j] + \log p(\rho_j \mid \tau) \geq \sum_{n=1}^{N} \sum_{k=1}^{K} \sum_{m=1}^{C_n^{(\text{major})}} q(z_n = k, U_n = j, M_n = m)$$
$$\times \Big[ \log \rho_j \{\Psi(B_n + \hat{\mu}^{\rho}_{n,j,m}) - \Psi(\hat{\mu}^{\rho}_{n,j,m}) + \Psi(D_n - B_n + \hat{\nu}^{\rho}_{n,j,m}) - \Psi(\hat{\nu}^{\rho}_{n,j,m})\}$$
$$- \rho_j \{\Psi(D_n + \hat{\rho}_j) - \Psi(\hat{\rho}_j)\}\Big] + \log \tau - \tau \rho_j + (\text{const.}) \equiv G_j(\rho_j \mid \tau).$$

Solve $\partial G_j(\rho_j \mid \tau)/\partial \rho_j = 0$:

$$\sum_{n=1}^{N}\sum_{k=1}^{K}\sum_{m=1}^{C_n^{(\mathrm{major})}} q(z_n = k, U_n = j, M_n = m)$$
$$\times \left[ \frac{\Psi(B_n + \hat{\mu}^{\rho}_{n,j,m}) - \Psi(\hat{\mu}^{\rho}_{n,j,m}) + \Psi(D_n - B_n + \hat{\nu}^{\rho}_{n,j,m}) - \Psi(\hat{\nu}^{\rho}_{n,j,m})}{\rho_j} - \{\Psi(D_n + \hat{\rho}_j) - \Psi(\hat{\rho}_j)\} \right] - \tau = 0$$
$$\Leftrightarrow \rho_j = \frac{\sum_{n=1}^{N}\sum_{m=1}^{C_n^{(\mathrm{major})}} \{\sum_{k=1}^{K} q(z_n = k, U_n = j, M_n = m)\}(\Xi^{\rho}_{n,j,m} + \Phi^{\rho}_{n,j,m})}{\tau + \sum_{n=1}^{N}\{\sum_{k=1}^{K}\sum_{m=1}^{C_n^{(\mathrm{major})}} q(z_n = k, U_n = j, M_n = m)\}\{\Psi(D_n + \hat{\rho}_j) - \Psi(\hat{\rho}_j)\}}, \quad \text{(S11)}$$

where

$$\Xi^{\rho}_{n,j,m} = \hat{\mu}^{\rho}_{n,j,m}\{\Psi(B_n + \hat{\mu}^{\rho}_{n,j,m}) - \Psi(\hat{\mu}^{\rho}_{n,j,m})\},$$
$$\Phi^{\rho}_{n,j,m} = \hat{\nu}^{\rho}_{n,j,m}\{\Psi(D_n - B_n + \hat{\nu}^{\rho}_{n,j,m}) - \Psi(\hat{\nu}^{\rho}_{n,j,m})\}.$$

In addition, $\tau$ is updated to the maximum likelihood as follows:

$$\tau = \frac{J}{\sum_{j=1}^{J} \rho_j}. \quad \text{(S12)}$$

#### 2 The detailed procedure to produce artificial mutation profiles

Here, we introduce how to establish artificial mutation profiles used in the simulation experiments. All datasets were produced through the generative process of SigTracer described in the main text. The parameters that were different for each dataset are shown in Table 1 in the main text, and here, we describe the common hyper-parameters and constants.

- Hyper-parameters of clone size $\pi$ : $\boldsymbol{\alpha}^{(\pi)} = \{\alpha_j^{(\pi)}\}_{j=1}^{J}$
  - $\alpha_j^{(\pi)}$ is set to 1.0 for all $j$; $1 \leq j \leq J$.
- Hyper-parameters of the signature activity of the $j$-th clone: $\boldsymbol{\alpha}_j^{(\theta)} = \{\alpha_{j,k}^{(\theta)}\}_{k=1}^{K}$
  - $\alpha_{j,k}^{(\theta)}$ is set to 1.0 for all $j$ and $k$; $1 \leq k \leq K$.
- Purity for each sample: $P$.
  - $\log(P)$ is derived from $\log(P) \sim \text{Uniform}(-1.0, 0.0)$, and obtain $P$ with exponential function.
- Copy number of the locus for mutation $n$ in normal cells: $C_n^{(\text{normal})}$.
  - $C_n^{(\text{normal})}$ is set to 2 for all $n$; $1 \leq n \leq N$.
- Copy number of the locus for mutation $n$ in tumor cells: $C_n^{(\text{tumor})}$.
  - For all $n$, $C_n^{(\text{tumor})}$ is derived from log-normal distribution with parameters $\mu = 1.0$ and $\sigma = 0.3$, whose probability density function is given as follows:

    $$p(c; \mu, \sigma) = \frac{1}{c\sigma\sqrt{2\pi}} \exp\left(-\frac{(\ln c - \mu)^2}{2\sigma^2}\right).$$

    Making the temporary variable $c$ an integer by a floor function, $C_n^{(\text{tumor})}$ is set to $\lfloor c \rfloor$ if $c \geq 1$. Otherwise, $C_n^{(\text{tumor})}$ is set to 1.
- Copy number of the locus for mutation $n$ in major alleles: $C_n^{(\text{major})}$.
  - For all $n$, the temporary variable $a$ is derived from the Beta distribution with shape parameters $\alpha = 5.0$ and $\beta = 3.0$. Then, $C_n^{(\text{major})}$ is determined by $\max(\lfloor C_n^{(\text{tumor})} \times a \rfloor, C_n^{(\text{tumor})} - \lfloor C_n^{(\text{tumor})} \times a \rfloor)$.

All samplings in the generation process was implemented using the numpy.random module in Python.

### 3 Supplementary results of the simulation experiment

**Table S2.** The predicted number of clones with artificial mutation profiles for the remaining datasets

| ID | Method | $J = 1$ | $J = 2$ | $J = 3$ | $J = 4$ | $J = 5$ |
|---|---|---|---|---|---|---|
| WES-2 | SigTracer | 13 | **86** | 1 | 0 | 0 |
| | CloneSig | 3 | **13** | 36 | 24 | 4 |
| WES-3 | SigTracer | 36 | **62** | 2 | 0 | 0 |
| | CloneSig | 2 | **17** | 33 | 21 | 27 |
| WES-4 | SigTracer | 49 | **51** | 0 | 0 | 0 |
| | CloneSig | 6 | **24** | 32 | 20 | 18 |
| WES-5 | SigTracer | 13 | **87** | 0 | 0 | 0 |
| | CloneSig | 0 | **33** | 24 | 19 | 24 |

Columns with $J = 1 \sim 5$ indicate the estimated number of signatures. Of the 100 samples in each dataset, the bold characters indicate the number of samples for which the correct number of clones could be estimated by each method. See Table 1 in the main manuscript for features of the datasets.

(**a**)

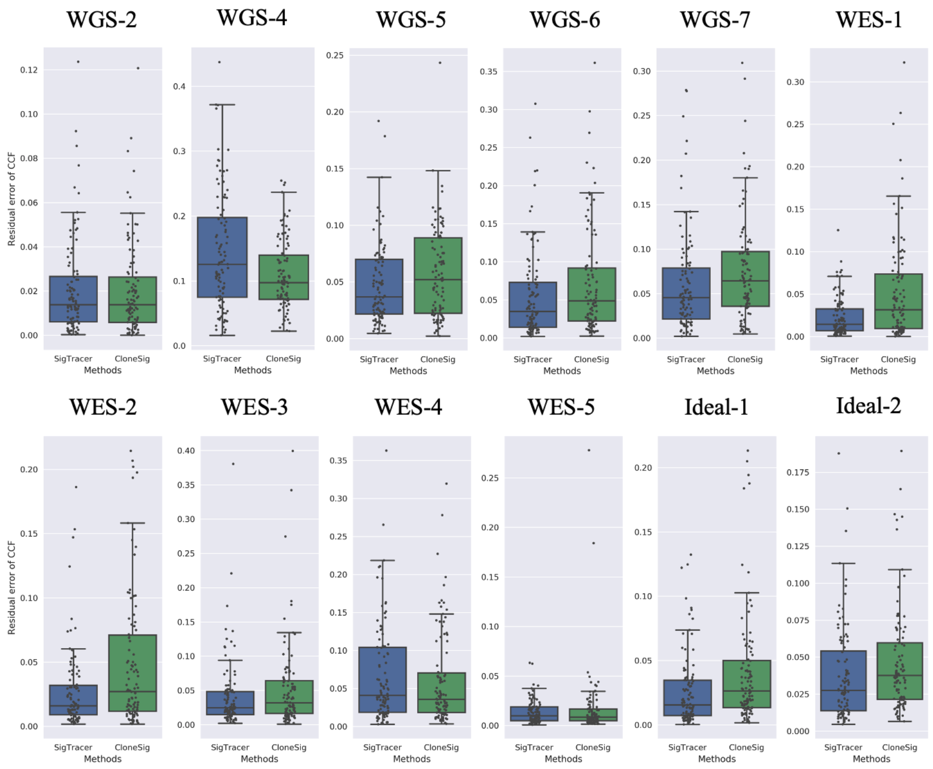


(**b**)

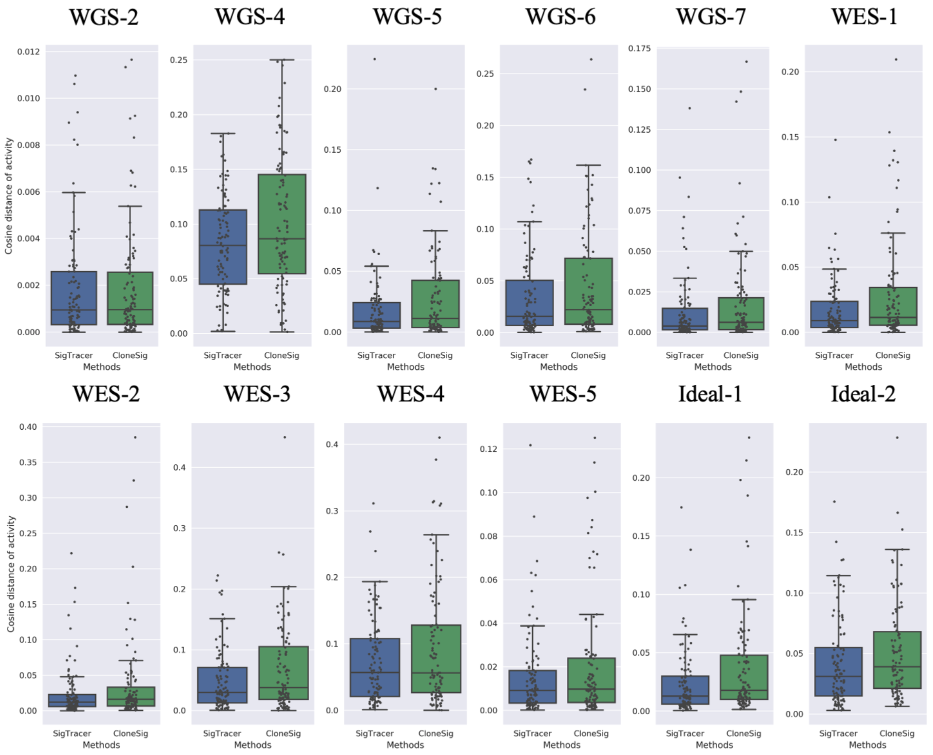


**Figure S1.** Accuracy of estimation for CCF and signature activity for the remaining datasets. (**a**) and (**b**) show the error of the estimated CCF and signature activity against the ground-truth values. Blue box plots are the results of SigTracer, and green box plots are those of CloneSig.

**Table S3.** Error between the true and estimated CCF in the simulation

| ID | Method | Mean (stdev) | Median(Q50) | Q25 | Q75 |
|---|---|---|---|---|---|
| WGS-1 | SigTracer | **0.064 (0.054)** | **0.044** | **0.027** | **0.084** |
| | CloneSig | 0.083 (0.065) | 0.071 | 0.036 | 0.108 |
| WGS-2 | SigTracer | 0.021 (0.022) | **0.014** | 0.006 | 0.027 |
| | CloneSig | **0.021 (0.021)** | 0.014 | **0.006** | **0.027** |
| WGS-3 | SigTracer | **0.104 (0.072)** | **0.087** | **0.047** | **0.148** |
| | CloneSig | 0.110 (0.072) | 0.094 | 0.056 | 0.152 |
| WGS-4 | SigTracer | 0.144 (0.091) | 0.126 | 0.075 | 0.199 |
| | CloneSig | **0.110 (0.054)** | **0.098** | **0.072** | **0.141** |
| WGS-5 | SigTracer | **0.048 (0.035)** | **0.037** | **0.021** | **0.070** |
| | CloneSig | 0.057 (0.004) | 0.052 | 0.022 | 0.089 |
| WGS-6 | SigTracer | **0.057 (0.059)** | **0.035** | **0.014** | **0.074** |
| | CloneSig | 0.072 (0.070) | 0.049 | 0.022 | 0.093 |
| WGS-7 | SigTracer | **0.061 (0.057)** | **0.046** | **0.022** | **0.080** |
| | CloneSig | 0.076 (0.060) | 0.065 | 0.036 | 0.098 |
| WES-1 | SigTracer | **0.024 (0.023)** | **0.015** | **0.007** | **0.033** |
| | CloneSig | 0.053 (0.061) | 0.032 | 0.009 | 0.075 |
| WES-2 | SigTracer | **0.027 (0.032)** | **0.016** | **0.009** | **0.032** |
| | CloneSig | 0.048 (0.052) | 0.027 | 0.012 | 0.071 |
| WES-3 | SigTracer | **0.043 (0.052)** | **0.025** | **0.015** | **0.049** |
| | CloneSig | 0.053 (0.064) | 0.032 | 0.016 | 0.065 |
| WES-4 | SigTracer | 0.068 (0.065) | 0.041 | 0.019 | 0.104 |
| | CloneSig | **0.061 (0.062)** | **0.036** | **0.019** | **0.071** |
| WES-5 | SigTracer | **0.014 (0.012)** | 0.010 | 0.005 | 0.019 |
| | CloneSig | 0.017 (0.033) | **0.009** | **0.005** | **0.017** |
| Ideal-1 | SigTracer | **0.027 (0.028)** | **0.016** | **0.007** | **0.035** |
| | CloneSig | 0.041 (0.045) | 0.027 | 0.013 | 0.050 |
| Ideal-2 | SigTracer | **0.039 (0.033)** | **0.028** | **0.014** | **0.055** |
| | CloneSig | 0.047 (0.035) | 0.038 | 0.022 | 0.060 |

The table shows the error between the true and estimated CCF in the simulation experiments. All values are desired to be small. Detailed information regarding each dataset is shown in Table 1 in the main paper. All statistical values were calculated for 100 artificial samples in each dataset. "stdev" denotes the standard deviation, and Q25(Q75) shows the average of the 25th and 26th (75th and 76th) smallest values out of 100 samples. In the comparison of the results between SigTracer and CloneSig, the smaller values are emphasized with bold text.

**Table S4.** Cosine distances between the true and estimated signature activity by each clone in the simulation experiments

| ID | Method | Mean (stdev) | Median(Q50) | Q25 | Q75 |
|---|---|---|---|---|---|
| WGS-1 | SigTracer | **0.023 (0.035)** | **0.011** | **0.003** | **0.028** |
| | CloneSig | 0.028 (0.034) | 0.012 | 0.003 | 0.038 |
| WGS-2 | SigTracer | **0.002 (0.002)** | **0.001** | **0.000** | **0.003** |
| | CloneSig | 0.002 (0.002) | 0.001 | 0.000 | 0.003 |
| WGS-3 | SigTracer | **0.058 (0.044)** | **0.045** | **0.024** | **0.087** |
| | CloneSig | 0.084 (0.068) | 0.070 | 0.031 | 0.121 |
| WGS-4 | SigTracer | **0.082 (0.044)** | **0.080** | **0.045** | **0.113** |
| | CloneSig | 0.101 (0.061) | 0.087 | 0.054 | 0.146 |
| WGS-5 | SigTracer | **0.018 (0.028)** | **0.009** | **0.003** | **0.025** |
| | CloneSig | 0.028 (0.037) | 0.011 | 0.004 | 0.043 |
| WGS-6 | SigTracer | **0.035 (0.041)** | **0.016** | **0.007** | **0.052** |
| | CloneSig | 0.045 (0.052) | 0.022 | 0.008 | 0.072 |
| WGS-7 | SigTracer | **0.013 (0.022)** | **0.004** | **0.002** | **0.015** |
| | CloneSig | 0.018 (0.030) | 0.006 | 0.002 | 0.022 |
| WES-1 | SigTracer | **0.018 (0.023)** | **0.009** | **0.004** | **0.024** |
| | CloneSig | 0.029 (0.039) | 0.011 | 0.006 | 0.035 |
| WES-2 | SigTracer | **0.025 (0.037)** | **0.012** | **0.006** | **0.023** |
| | CloneSig | 0.037 (0.063) | 0.016 | 0.007 | 0.035 |
| WES-3 | SigTracer | **0.049 (0.051)** | **0.030** | **0.013** | **0.071** |
| | CloneSig | 0.069 (0.073) | 0.038 | 0.018 | 0.107 |
| WES-4 | SigTracer | **0.074 (0.064)** | 0.057 | **0.021** | **0.108** |
| | CloneSig | 0.095 (0.091) | **0.056** | 0.026 | 0.128 |
| WES-5 | SigTracer | **0.015 (0.019)** | **0.009** | **0.003** | **0.018** |
| | CloneSig | 0.02 (0.027) | 0.010 | 0.003 | 0.024 |
| Ideal-1 | SigTracer | **0.024 (0.029)** | **0.013** | **0.006** | **0.030** |
| | CloneSig | 0.037 (0.046) | 0.018 | 0.010 | 0.048 |
| Ideal-2 | SigTracer | **0.042 (0.036)** | **0.031** | **0.015** | **0.055** |
| | CloneSig | 0.052 (0.041) | 0.039 | 0.021 | 0.068 |

The table shows the cosine distance between the true and estimated signature activity by each clone in the simulation experiments. This table is similar to Table S3.

#### 4 Derivation of the method of calculating the reconstruction rate and the result for each dataset

Denoting $M_v$ as the proportion of $hv$-th type mutations in a sample, then $RR$ for mutation , $RR_{\text{mutation}}$ was defined as follows:

$$RR_{\text{mutation}} \equiv \frac{1}{V}\sum_{v=1}^{V} \min(M_v, \hat{M}_v)$$
$$\text{with } \hat{M}_v = \sum_{k=1}^{K}\left\{\sum_{j=1}^{J}\pi_j\theta_{j,k}\right\}\phi_{k,v}.$$

$RR$ for VAF was complicated because VAF assumed a continuous value. Therefore, we defined $RR$ for VAF, $RR_{\text{VAF}}$, by dividing the value range of the observed VAF and the VAF distribution based on the estimated parameters (a beta mixture) into 30 partitions and comparing the expected number of mutations included in each partition. Denoting $\varepsilon_i$ as the number of mutations included in the $i$-th partition and the segment of the $i$-th partition $[s_i^{\text{start}}, s_i^{\text{end}})$ holding $s_i^{\text{end}} = s_{i+1}^{\text{start}}$, $RR_{\text{VAF}}$ was defined as follows:

$$RR_{\text{VAF}} \equiv \frac{1}{N}\sum_{i=1}^{30}\min(\varepsilon_i, \hat{\varepsilon}_i)$$
$$\text{with } \hat{\varepsilon}_i = \sum_{n=1}^{N}\sum_{j=1}^{J}\pi_j\int_{s_i^{\text{start}}}^{s_i^{\text{end}}}\text{Beta}(s \mid \mu_{n,j}, \nu_{n,j})ds$$

where $\text{Beta}(s \mid \mu, \nu)$ is a probability density function of the beta distribution with random variable $s$ according to mean $\mu$ and overdispersion $\nu$. We calculated the integral explicitly using the regularized incomplete beta function.

**Table S5.** Reconstruction rate for each dataset

| ID | $RR_{\text{mutation}}$ Mean (stdev) | $RR_{\text{mutation}}$ Median | $RR_{\text{VAF}}$ Mean (stdev) | $RR_{\text{VAF}}$ Median |
|---|---|---|---|---|
| WGS-1 | 0.933 (0.009) | 0.931 | 0.718 (0.078) | 0.715 |
| WGS-2 | 0.936 (0.010) | 0.935 | 0.819 (0.048) | 0.831 |
| WGS-3 | 0.932 (0.008) | 0.932 | 0.641 (0.063) | 0.641 |
| WGS-4 | 0.934 (0.009) | 0.934 | 0.626 (0.066) | 0.627 |
| WGS-5 | 0.934 (0.010) | 0.934 | 0.695 (0.081) | 0.692 |
| WGS-6 | 0.933 (0.009) | 0.933 | 0.709 (0.078) | 0.717 |
| WGS-7 | 0.936 (0.011) | 0.935 | 0.711 (0.077) | 0.709 |
| WES-1 | 0.906 (0.013) | 0.906 | 0.800 (0.100) | 0.807 |
| WES-2 | 0.868 (0.019) | 0.865 | 0.807 (0.088) | 0.818 |
| WES-3 | 0.787 (0.032) | 0.786 | 0.804 (0.089) | 0.832 |
| WES-4 | 0.709 (0.048) | 0.706 | 0.797 (0.075) | 0.809 |
| WES-5 | 0.905 (0.015) | 0.906 | 0.807 (0.104) | 0.838 |
| Ideal-1 | 0.934 (0.008) | 0.935 | 0.786 (0.071) | 0.800 |
| Ideal-2 | 0.932 (0.009) | 0.932 | 0.730 (0.066) | 0.723 |
| CLL | 0.844 (0.025) | 0.843 | 0.669 (0.049) | 0.664 |
| BNHL | 0.865 (0.024) | 0.865 | 0.678 (0.070) | 0.682 |

The table shows the $RR$ values calculated from the results of all experiments by SigTracer. The statistical values for all samples included in each dataset are listed. "stdev" denotes the standard deviation.

#### 5 Supplementary result with real mutation profiles from the PCAWG cohort

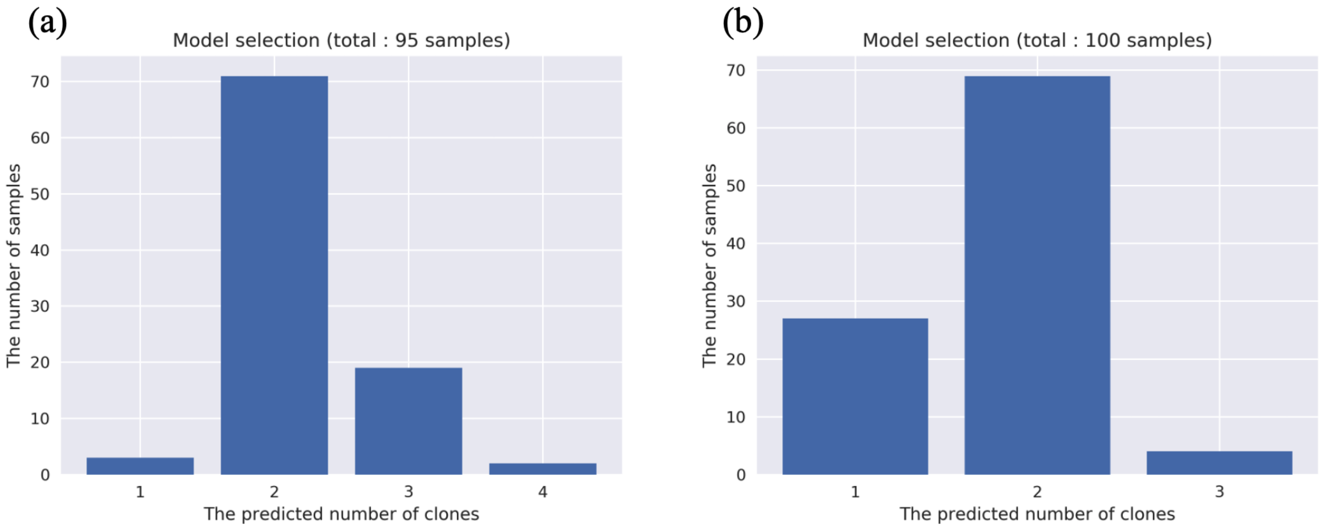


**Figure S2.** The predicted number of clones in the CLL and BNHL samples. (**a**) and (**b**) show the predicted number of clones in CLL and BNHL samples, respectively.

**Table S6.** Frequently mutated region in clones with high activity of SBS1, SBS5, and SBS40 for CLL samples

**SBS1**

| Hugo symbol | variant classification | p-value $<$ FWER |
|---|---|---|
| AC018717.1 | lincRNA | 6.64E-08 |

**SBS5**

| Hugo symbol | variant classification | p-value $<$ FWER |
|---|---|---|
| IGLL5 | Intron | 1.18E-06 |

**SBS40**

| Hugo symbol | variant classification | p-value $<$ FWER |
|---|---|---|
| AC018717.1 | lincRNA | 8.37E-10 |
| IGLV2-14 | RNA | 5.136E-07 |

This table lists the mutated regions significantly correlated with SBS1, SBS5, and SBS40 in CLL samples detected according to the test pipeline described in the main text. Only those that were determined to be significant by Bonferroni correction are shown.

**Table S7.** Frequently mutated regions in clones with high SBS9 activity for BNHL samples

| Hugo symbol | variant classification | p-value < FWER |
|---|---|---|
| FHIT | Intron | 1.73E-46 |
| Unknown | IGR | 4.72E-34 |
| PCDH15 | Intron | 4.67E-21 |
| CADM2 | Intron | 2.58E-18 |
| ROBO2 | Intron | 9.31E-17 |
| LPHN3 | Intron | 1.87E-16 |
| LAPTM5 | Intron | 2.02E-15 |
| GHR | Intron | 1.26E-14 |
| CCSER1 | Intron | 4.98E-13 |
| RP11-6N13.1 | lincRNA | 5.24E-13 |
| ROBO1 | Intron | 7.07E-13 |
| CSMD3 | Intron | 1.00E-11 |
| GRID2 | Intron | 3.14E-11 |
| DISC1FP1 | RNA | 3.48E-11 |
| GRIA2 | Intron | 1.37E-10 |
| PCDH10 | Intron | 1.41E-10 |
| CTC-340A15.2 | RNA | 4.13E-09 |
| CDH18 | Intron | 6.01E-09 |
| LRP1B | Intron | 2.63E-08 |
| IGLV3-22 | RNA | 9.09E-08 |
| LPP | Intron | 9.72E-08 |
| CTC-535M15.2 | lincRNA | 1.57E-07 |
| RP11-427M20.1 | lincRNA | 1.97E-07 |
| DGKB | Intron | 2.29E-07 |
| SEMA3A | Intron | 2.57E-07 |
| NAALADL2 | Intron | 6.95E-07 |
| AC096579.13 | RNA | 7.87E-07 |
| LINC00669 | lincRNA | 8.19E-07 |

The table lists the mutated regions significantly correlated with SBS9 in BNHL samples detected according to the test pipeline described in the main text. Only those mutations that were determined to be significant by Bonferroni correction are shown.

**Table S8.** The number of significantly mutated regions with a simple significance level, $\alpha < 0.05$.

| Signature | CLL | BNHL | Common |
|---|---|---|---|
| SBS1 | 126 | 228 | 9 |
| SBS5 | 84 | 222 | 13 |
| SBS9 | 308 | 766 | 74 |
| SBS40 | 150 | 536 | 17 |

There were 23046 and 52538 annotations in CLL and BNHL datasets, respectively. This table shows the number of significantly mutated regions in the clones with high activity of each signature among them.

**Table S9.** Significantly mutated regions in both CLL and BNHL samples

| Signature | Mutated region |
|---|---|
| SBS1 | CTNNA2(Intron), CXCR4(Intron), EBF1(Intron), ELOVL7(Intron), FHIT(Intron), FSTL5(Intron), IGHM(RNA), SYNPR(Intron), ZCCHC7(Intron), |
| SBS5 | BCL6(Intron), CTNNA2(Intron), EBF1(Intron), FHIT(Intron), FSTL5(Intron), GRIA2(Intron), IGHM(RNA), IMMP2L(Intron), KIAA0125(5'Flank), PAX5(Intron), RBFOX1(Intron), RP1-209A6.1(lincRNA), ZCCHC7(Intron), |
| SBS9 | AC006373.1(lincRNA), AC092661.1(lincRNA), AC096579.13(RNA), CADM2(Intron), CDH12(Intron), CDH18(Intron), CDH19(Intron), CNTN5(Intron), CNTNAP5(Intron), CSMD3(Intron), CTC-340A15.2(RNA), CTC-535M15.2(lincRNA), CTD-2134P3.1(lincRNA), CTD-2374C24.1(lincRNA), CTNNA2(Intron), DGKB(Intron), DISC1FP1(RNA), DPYD(Intron), EPHA7(Intron), FHIT(Intron), FSTL5(Intron), GRID2(Intron), GRM5(Intron), GRM7(Intron), IGHV1-8(RNA), IGHV4-34(RNA), IGKV4-1(RNA), IGLV3-25(RNA), IGLV7-46(RNA), IMMP2L(Intron), KLHL6(Missense Mutation), LINC00669(lincRNA), LINC01036(lincRNA), LPHN3(Intron), LPP(Intron), LRP1B(Intron), LUZP2(Intron), NEGR1(Intron), NKAIN2(Intron), OGDH(Intron), PCDH15(Intron), PCDH9(Intron), PCLO(Intron), ROBO2(Intron), RP11-138I17.1(lincRNA), RP11-149A7.2(lincRNA), RP11-184E9.1(lincRNA), RP11-389E17.1(Intron), RP11-398M15.1(lincRNA), RP11-3L8.3(RNA), RP11-400D2.3(lincRNA), RP11-404I7.2(lincRNA), RP11-427M20.1(lincRNA), RP11-44H4.1(lincRNA), RP11-453M23.1(lincRNA), RP11-506N2.1(lincRNA), RP11-553P9.2(lincRNA), RP11-659O3.1(lincRNA), RP11-6N13.1(lincRNA), RP11-734I18.1(lincRNA), RP11-74C3.1(lincRNA), RP11-79E3.3(lincRNA), RP3-463P15.1(lincRNA), RP5-887A10.1(lincRNA), SEMA3A(Intron), SEMA3C(Intron), SGCZ(Intron), SLC5A8(Intron), SNTG1(Intron), TCL1A(Intron), TMTC2(Intron), TUSC3(Intron), Unknown(IGR), ZFPM2(Intron), |
| SBS40 | CADM2(Intron), CDH10(Intron), DENND1B(Intron), DLEU1(Intron), ESR2(Intron), F11-AS1(RNA), IGLL5(Intron), IGLV2-14(RNA), PACSIN1(Intron), PAX5(Intron), RP11-184E9.1(lincRNA), RP11-463J7.2(lincRNA), RP11-638L3.1(lincRNA), RP11-89M20.2(lincRNA), SIPA1L1(Intron), STAU2(Intron), ZFP36L1(Intron), |

It shows significantly mutated regions in both CLL and BNHL samples by each signature. Each symbol, "A(B)", indicates that the variant classification is B for the gene annotated as A in Hugo symbol.

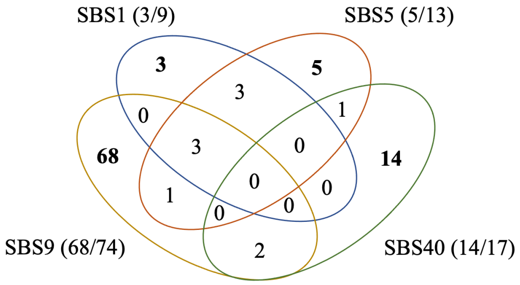


**Figure S3.** Unique mutated regions with respect to the signature. The Venn diagram shows whether the mutated regions shown in Table S9 overlap between signatures. In particular, we can see that most mutated regions of SBS9 are unique.
